# A RID-like cytosine methyltransferase homologue controls sexual development in the fungus *Podospora anserina*

**DOI:** 10.1101/575746

**Authors:** P Grognet, H Timpano, F Carlier, J Aït-Benkhali, V Berteaux-Lecellier, R Debuchy, F Bidard, F Malagnac

## Abstract

DNA methyltransferases are ubiquitous enzymes conserved in bacteria, plants and opisthokonta. These enzymes, which methylate cytosines, are involved in numerous biological processes, notably development. In mammals and higher plants, methylation patterns established and maintained by the cytosine DNA methyltransferases (DMTs) are essential to zygotic development. In fungi, some members of an extensively conserved fungal-specific DNA methyltransferase class are both mediators of the Repeat Induced Point mutation (RIP) genome defense system and key players of sexual reproduction. Yet, no DNA methyltransferase activity of these purified RID (RIP deficient) proteins could be detected *in vitro*. These observations led us to explore how RID-like DNA methyltransferase encoding genes would play a role during sexual development of fungi showing very little genomic DNA methylation, if any.

To do so, we used the model ascomycete fungus *P. anserina*. We identified the *PaRid* gene, encoding a RID-like DNA methyltransferase and constructed knocked-out Δ*PaRid* defective mutants. Crosses involving *P. anserina* Δ*PaRid* mutants are sterile. Our results show that, although gametes are readily formed and fertilization occurs in a Δ*PaRid* background, sexual development is blocked just before the individualization of the dikaryotic cells leading to meiocytes. Complementation of Δ*PaRid* mutants with ectopic alleles of *PaRid*, including GFP-tagged, point-mutated, inter-specific and chimeric alleles, demonstrated that the catalytic motif of the putative PaRid methyltransferase is essential to ensure proper sexual development and that the expression of PaRid is spatially and temporally restricted. A transcriptomic analysis performed on mutant crosses revealed an overlap of the PaRid-controlled genetic network with the well-known mating-types gene developmental pathway common to an important group of fungi, the Pezizomycotina.

**Author Summary:** Sexual reproduction is considered to be essential for long-term persistence of eukaryotic species. Sexual reproduction is controlled by strict mechanisms governing which haploids can fuse (mating) and which developmental paths the resulting zygote will follow. In mammals, differential genomic DNA methylation patterns of parental gametes, known as ‘DNA methylation imprints’ are essential to zygotic development, while in plants, global genomic demethylation often results in female-sterility. Although animal and fungi are evolutionary related, little is known about epigenetic regulation of gene expression and development in multicellular fungi. Here, we report on a gene of the model fungus *Podospora anserina*, encoding a protein called PaRid that looks like a DNA methyltrasferase. We showed that expression of the catalytically functional version of the PaRid protein is required in the maternal parental strain to form zygotes. By establishing the transcriptional profile of *PaRid* mutant strain, we identified a set of PaRid direct and/or indirect target genes. Half of them are also targets of a mating-type transcription factor known to be a major regulator of sexual development. So far, there was no other example of identified RID targets shared with a well-known developmental pathway that is common to an important group of fungi, the Pezizomycotina

## Introduction

A covalently modified DNA base, the 5-methylcytosine (5-meC) is common in genomes of organisms as diverse as bacteria, fungi, plants and animals. In eukaryotes, when present, this epigenetic modification is associated with down-regulation of gene expression and suppression of transposon activity. Patterns of cytosine methylation are established and maintained through subsequent DNA replication cycles by the cytosine DNA methyltransferases (DMTs), a class of enzymes conserved from bacteria to mammals. Five families of eukaryotic DMTs can be distinguished [1,2]: 1) the “maintenance” DMT family which includes the mammalian DNMT1 [3], the plant MET1 [4] and the fungal DIM-2 enzymes [5]; 2) the “*de novo*” DMT family which includes the mammalian DNMT3A and DNMT3B [6] and the plant Domains Rearranged Methyltransferase DRM2 [7]; 3) the flowering plant-specific “maintenance” chromomethylase (CMT) family, which includes the *Arabidopsis thaliana* CMT2 and CMT3 enzymes [8–10]; 4) the fungal-specific DMT-like family which includes the *Ascobolus immersus* Masc1 [11] and the *Neurospora crassa* RID proteins [12]; 5) the putative CG-specific “maintenance” DMTs of the structurally divergent DNMT5 family [13,14]. DMTs from the first three families have been shown to methylate cytosines *in vitro*, while no such activity has ever be demonstrated either for the fungal DMT-like protein [11,12] nor for the DMTs of the DNMT5 family [13]. A sixth family, typified by Dnmt2 which is now known to be a tRNA methyltransferase, was therefore discarded from the *bona fide* DMT group [15]. Surprisingly, genes encoding putative DMTs can also be found in genome of species actually having very few, if any, 5-meC (*e.g., Dictyostelium discoideum* [16], *Drosophila melanogaster* [17] and *Aspergillus nidulans* [18]). These DMT-like proteins might be endowed with a still undetermined function. By contrast, most of the ascomycetous yeasts, including the model organism *Saccharomyces cerevisiae* and the human pathogen *Candida albicans*, lack genes encoding putative DMT-like enzymes. A notable exception includes the fission yeast *Schizosaccharomyces pombe* genome that contains a *Dnmt2*-like homolog. To date, the presence of 5-mC in the ascomycetous yeast genomes remains controversial [19,20].

In mammals and higher plants, methylation of cytosines is mostly associated with stable and inheritable repression of gene transcription. Therefore, it plays an essential role in genome stability and developmental programs such as imprinting and X chromosome inactivation [21]. In mice, loss of function of DNMT1 results in embryonic lethality [22] whereas flowering plants that present mutation in MET1, display a spectrum of viable developmental abnormalities [23,24]. In fungi, however, alteration of 5-meC content results in more contrasted outcomes. In the model ascomycete *N. crassa* inactivation of the *dim-2* gene, which encodes an enzyme from the “maintenance” DMT family results in complete loss of DNA methylation. Remarkably, the *dim-2* mutants behave as wild-type strains with respect to their vegetative life cycle and their sexual reproduction cycle [5]. These observations indicate that although *N. crassa* displays substantial cytosine methylation in some genomic compartments, this epigenetic modification is dispensable for all developmental processes observed in laboratory conditions. RID (for RIP Deficient), the second putative DMT of *N. crassa* belongs to the fungal-specific DMT-like family. The RID protein plays an essential role in RIP (Repeat-Induced point mutation), a genome defense mechanism conserved among Pezizomycotina fungi [12,25,26]. As defined in *N. crassa* by Selker and colleagues, RIP occurs in haploid parental nuclei after fertilization but before karyogamy. Repetitive DNA sequences originally described as longer than 400 bp, are detected during the RIP process and are subsequently subjected to extensive conversion of cytosine to thymidine (C-to-T) [27,28]. Over the extent of the RIP targeted repeats, the remaining cytosines are heavily methylated. Furthermore, in *N. crassa*, recent findings showed that linker regions located in between RIPed repeats are subjected to RID-independent RIP [29]. This cytosine to thymine mutagenic process is mediated by DIM-2 and relies on heterochromatin-related pathway. In this fungus, the absence of RID abolishes RIP on newly formed repeats without causing additional defect, such mutants displaying wild-type vegetative growth and sexual cycle.

The analogous MIP (Methylation Induced Premeiotically) process discovered in *A. immersus* leads to cytosine methylation only, within repeats [30]. Disruption of *Masc1*, which encodes a member of the fungal-specific DMT-like family, results in abolition of the *de novo* methylation of repeats but also in severely impaired sexual development [11]. Indeed, when *Masc1* is absent in both parental stains, the resulting crosses are arrested at an early stage of sexual reproduction and no dikaryotic cells formed. Similar defects of sexual development have been observed when RID-like *DmtA* and *TrRid* knockout mutants were constructed in *A. nidulans* [18] and *Trichoderma reesei* [31], respectively (Wan-Chen Li and Ting-Fang Wang, personal communication). This suggests that, unlike RID of *N. crassa*, several members of the fungal-specific DMT-like family could be involved in both genome defense and development [32], as for Masc1, which appears as the founding prototype of this family of putative DMTs.

Unlike *A. nidulans* and *T. reesei*, the model ascomycete, *P. anserina* do not produce asexual conidia and therefore relies only on sexual reproduction to propagate, before the colony dies of senescence. Besides, this fungus displays an active RIP process prior to the formation of the zygotes, but the RIPed targets are not methylated [33,34]. In this context, we explored the function of PaRid, a DMT homologue closely related to the *N. crassa* RID protein. In the present study, we showed that the PaRid fungal-specific DMT-like protein is essential to ensure proper sexual development, while the catalytically dead version of this protein could not. Moreover, we demonstrated that PaRid is required in the maternal parental strain. Indeed, in the absence of PaRid, even if the wild-type allele is present in the nuclei coming from the male gamete, the sexual development is blocked before the dikaryon formation. By establishing the transcriptional profile of mutant crosses, we also identified a set of PaRid direct and/or indirect target genes. A substantial subset of these genes was previously identified as targets of FPR1, a mating-type transcription factor known to be a major regulator of fertilization and subsequent sexual development.

## Results

### PaRid is essential to ensure proper sexual development

The *P. anserina* genome contains a single gene, namely *PaRid* (Pa_1_19440) [35], which encodes a 752 amino acid protein closely related to fungal-specific DMT-like proteins (Fig 1A). In addition of conserved DMT domain motifs (I to X) and expected Target Recognition Domain variable region (TRD), PaRid shows typical signatures of cytosine methyltransferases (PF00145, PS51679, PR00105) (Fig 1B). This protein belongs to the same phylogenetic group (DMT-like fungal-specific family) as the previously described *N. crassa* RID protein (41% identity) [18], *A. immersus* Masc1 protein (33% identity) [11] and *A. nidulans* DmtA (31% identity) (Fig 1C). As already pointed out for Masc1/Rid DMT-like, these proteins, including PaRid, have peculiar features [32]. They show a non-canonical EQT (glutamate-glutamine-threonine) triad in motif VI instead of the ENV (glutamate-asparagine-valine) triad shared by all other eukaryotic C5-cytosine methyltransferases (Fig 1D) as well as a remarkably short TRD domain (80 amino acids).

**Fig 1.**
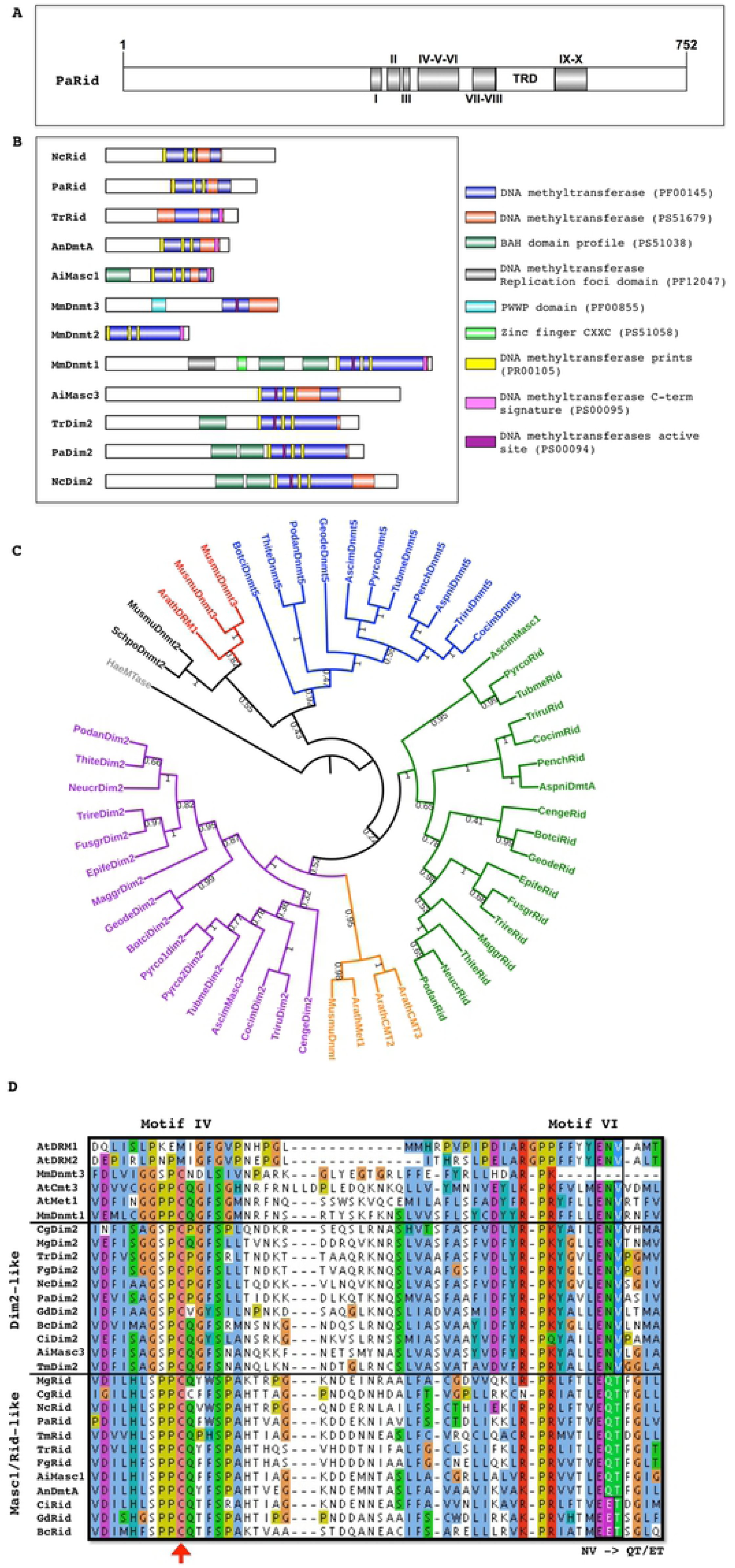
Structure and Phylogenetic analysis of DNA methyltransferases proteins. (**A**) Domain architecture of *P. anserina* putative DNA methyltransferase PaRid. The catalytic domains contain 10 conserved motifs (I - X) and a target recognition domain (TRD) located between the motifs VIII and IX. The amino acid length is indicated. (**B**) Domain architecture of DNA methyltransferase proteins. The functional domain analysis was performed using InterProScan and visualized using IBS. *Mus musculus*: MmDnmt1 (AAH53047.1), MmDnmt2 (AAC40130.1), MmDnmt3 (AAF73868.1); *Ascobolus immersus*, AiMasc1 (AAC49849.1), AiMasc3; *Aspergillus nidulans* AnDmtA (AAO37378.1), *Neurospora crassa* NcRid (AAM27415.1), NcDim2 (AAK49954.1); *Podospora anserina* PaRid, PaDim2; *Trichoderma reesei* TrRid (AEM66210.1), TrDim2 (XP_006964860.1). Cytosine-specific DNA methyltransferase domains: PF00145, PR00105, PS51679, PS00095, PS00094; protein-DNA interaction domains: bromo-associated homology (BAH) domain PS51038, Replication foci targeting sequence (RFTS) PF12047, Zinc finger motif (CXXC) PS51058, PWWP domain (PF00855). Because its sequence is too divergent, PaDnmt5 (Pa_4_2960) was not included. (**C**) Phylogenetic analysis of mammal, plant and fungal DNA methyltransferases. The maximum likelihood tree resolved five groups i) the Dnmt1/Met1/CMT group (orange), ii) the Dnmt2, group (black) iii) the Dnmt3/DRM group (red) iv) the fungal Dim2-like group (purple) and v) the fungal-specific Rid-like group (green), vi) the DNMT5 group (blue). *Haemophilus aegyptius* (HaeMTase: WP_006996493.1) *Arabidopsis thaliana* (ArathMet1: NP_199727.1; ArathCMT2: NP_193637.2; ArathCMT3: NP_177135.1; ArathDRM1: NP_197042.2; ArathDRM2: NP_196966.2), *Ascobolus immersus* (AscimMasc3: CE37440_11164; AscimMasc1: AAC49849.1; AscimDnmt5: RPA73956.1), *Aspergillus nidulans* (AspniDmtA: XP_664242.1, AspniDnmt5: XP_663680.1), *Botrytis cinerea* (BotciDim2: XP_024553164.1; BotciRid: XP_024550989.1, BotciDnmt5: XP_024550790.1), *Cenococcum geophilum* (CengeDim2: OCK96497.1; CengeRid: OCK89234.1), *Coccidioides immitis* (CocimDim2: XP_001247991.2; CocimRid: XP_001239116.2; CocimDnmt5: XP_001247253.2) *Epichloe festucae* (EpifeDim2: annotated in this study; EpifeRid: AGF87103.1), *Fusarium graminearum* (FusgrDim2: EYB34029.1; FusgrRid: XP_011320094.1) *Pseudogymnoascus destructans* (GeodeDim2: XP_024321957.1; GeodeRid: XP_024328520.1; GeodeDnmt5: XP_024320712.1) *Magnaporthe grisea* (MaggrDim2: XP_003718076.1; MaggrRid: XP_003720946.1) *Mus musculus* (MusmuDnmt1: NP_001300940.1; MusmuDnmt3A: NP_031898.1; MusmuDnmt3B: NP_001003961.2, MusmuDnmt2: NP_034197.3), *Neurospora crassa* (NeucrDim2 : XP_959891.1; NeucrRid : AAM27408.1), *Penicillium chrysogenum* (PenchRid: XP_002563814.1; PenchDnmt5: XP_002561360.1), *Podospora anserina* (PodanDim2: Pa_5_9100; PodanRid: Pa_1_19440; PodanDnmt5: Pa_4_2960), *Pyronema confluens* (Pyrco1dim2: PCON_02009m.01; Pyrco2dim2: PCON_01959m.01, PircoRid: PCON_06255m.01; PyrcoDnmt5: CCX08765.1), *Schizosaccharomyces pombe* (SchpoDnmt2 : NP_595687.1), *Thielavia terrestris* (ThiteDim2: XP_003654318.1; ThiteRid: XP_003651414.1; ThiteDnmt5: XP_003650845.1), *Trichoderma reesei* (TrireDim2 XP_006964860.1; TrireRid: AEM66210.1), *Trichophyton rubrum* (TriruDim2: XP_003239082.1; TriruRid: XP_003239287.1; TriruDnmt5: XP_003236242.1), *Tuber melanosporum* (TubmeDim2: XP_002837027.1; TubmeRid: XP_002842459.1; TubmeDnmt5: XP_002837747.1). (**D**) Comparison of the amino acid sequence of the catalytic motifs IV and VI. A key catalytic step is the nucleophilic attack of the DNA methyltransferases on the sixth carbon of the target cytosine. This attack is made by the cysteine residue (red arrow) of the conserved PCQ triad (motif IV). This reaction is catalyzed by protonation of the N3 position of the cytosine by the glutamate residue of the conserved ENV triad (motif VI). In the Masc1/Rid-like group of enzymes, the ENV triad is replaced by either the EQT triad (e.g. *N. crassa* Rid, *A. immersus* Masc1, *P. anserina* Rid, etc.) or the EET triad (e.g. *B. cinerea* Rid, *P. destructans* Rid, etc.). At: *Arabidopsis thaliana*, Mm: *Mus musculus*, Cg: *Cenococcum geophilum*, Mg: *Magnaporthe grisea*, Tr: *Trichoderma reesei*, Fg: *Fusarium graminearum*, Nc: *Neurospora crassa*, Pa: *Podospora anserina*, Gd: *Pseudogymnoascus destructans*, Bc: *Botrytis cinerea*, Ci: *Coccidioides immitis*. Ai: *Ascobolus immersus*, Tm: *Tuber melanosporum*, An: *Aspergillus nidulans*. See above for accession numbers of the corresponding proteins.

An EST database generated from vegetatively grown *P. anserina* showed no expression of *PaRid*. However, the *PaRid* expression profile extracted from a microarray transcriptional time-course analysis performed during *P. anserina*’s sexual development (Bidard and Berteaux-Lecellier, GEO accession no. GSE104632) showed a peak at T12 (12 hours post fertilization) followed by a decrease at T30 (30 hours post fertilization) (S1A Fig). Besides RT-PCR experiments (S1B Fig) indicated that *PaRid* transcripts could be detected up to T96 (96 hours post fertilization). Searching the available *P. anserina* RNA-seq data [36], we did not find any evidence of non-coding RNA at the *PaRid* locus (S1C Fig), as described for sense-antisense DmtA/tmdA transcripts found in *A. nidulans* [18].

To gain insight about a potential function of PaRid during the life cycle of *P. anserina*, we deleted the corresponding gene. Replacement of the *PaRid* wild-type allele with the hygromycin-B resistance marker generated the Δ*PaRid* null allele (*PaRid::hph*) (Materials and methods and S2 Fig). The Δ*PaRid* mutant strains displayed wild-type phenotypes with respect to vegetative growth and vegetative developmental programs (i.e. germination, mycelium pigmentation, aerial hyphae production, branching, anastomosis, longevity, stress resistance, See S2 Table, S4 and S5 Figs), indicating that the *PaRid* gene is dispensable for the vegetative phase of *P. anserina*’s life cycle.

We then investigated the ability of the Δ*PaRid* mutant strains to perform sexual reproduction (see S3 Fig for description). During the first 30 hours post-fertilization, the sexual development of homozygous Δ*PaRid* crosses is indistinguishable from that of wild-type crosses. In homozygous Δ*PaRid* crosses, the fruiting bodies are formed normally, both in timing (24 hours post-fertilization) and in number (4333 in average per cross +/- 368, N=5), as compared to homozygous wild-type crosses (4477 in average per cross +/- 458, N=5) (S3 Table, Fig 2A). However, while the fructifications originating from homologous wild-type crosses continued to develop up to 45 hours post-fertilization, those originating from homozygous Δ*PaRid* crosses stopped maturing around 30 hours after fertilization, forming distinctive micro-perithecia only. Although these micro-perithecia never grew into fully mature fruiting bodies, they displayed no visible morphological defects (Fig 2B). Importantly, they were fully pigmented and harbored typical necks and ostioles. As a consequence of this early developmental arrest, while one fully matured perithecium would produce hundreds of asci after 96 hours post fertilization, the Δ*PaRid* micro-perithecia are barren (Fig 2C).

**Fig 2.**
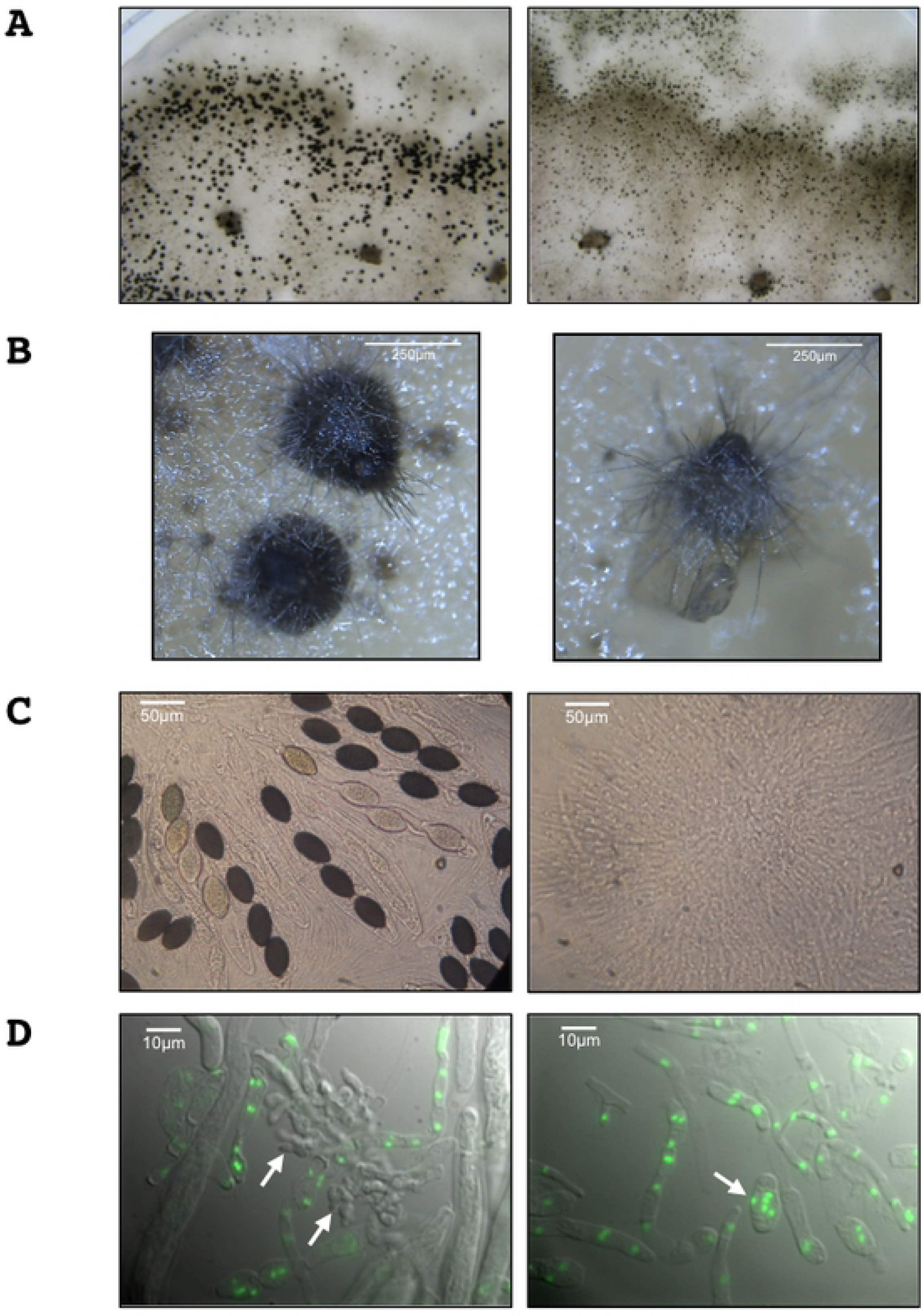
PaRid is essential to complete fruiting body development and to produce ascospores. (**A**) Homozygous crosses of wild type S strains (left panel) and of Δ*PaRid* strains (right panel) on M2 medium after 5 days at 27°C. Each dark dot is one fruiting body resulting from one event of fertilization. The homozygous Δ*PaRid* cross forms reduced-size fruiting bodies only (right panel). (**B**) Close up of fruiting bodies (perithecia) originating from either a wild-type genetic background (left panel) or a Δ*PaRid* genetic background (right panel). Scale bar: 250 µm. (**C**) After 4 days of growth at 27°C, the wild type fruiting bodies start to produce ascospores (left panel) while the mutant micro-perithecia are barren (right panel). Scale bar: 50 µm. (**D**) Fluorescence microscopy pictures of 48h-old fruiting body content from homozygous crosses of wild type S strains (left panel) and of Δ*PaRid* strains (right panel), performed on M2 medium at 27°C. The nuclei are visualized thanks to histone H1-GFP fusion protein. Croziers are readily formed inside the wild type perithecia (left panel, white arrows) while no crozier but large plurinucleate ascogonial cells only are seen inside the Δ*PaRid* perithecia. Scale bar: 10 µm.

Furthermore, heterozygous orientated crosses showed that when the wild-type *PaRid* allele was present in the female gamete genome (i.e. ascogonia) and the Δ*PaRid* null allele was present in the male gamete genome (i.e. spermatia), the fruiting-body development was complete and resulted in the production of asci with ascospores (S6B Fig). The progeny isolated from this cross showed the expected 1:1 segregation of the Δ*PaRid* allele. On the contrary, when the Δ*PaRid* null allele was present in the female gamete genome and the wild-type *PaRid* allele was present in the male gamete genome, the fruiting body development was blocked and the resulting micro-perithecia are barren (S6B Fig). Altogether, these results indicate that (1) Δ*PaRid* mutants formed both male and female gametes and that these gametes are able to fuse since (2) fertilization occurred as efficiently in homozygous Δ*PaRid* mutant crosses than in wild-type crosses, yielding a similar number of fruiting bodies per crosses (3) the wild-type *PaRid* allele must be present in the maternal haploid genome for completion of sexual development. Notably, because reciprocal heterozygous orientated crosses with *mat-* and *mat+* wild-type strains behaved similarly, the observed Δ*PaRid* phenotype was not mating-type dependent.

### PaRid does not contribute to the development of maternal tissues

Fruiting bodies in Pezizomycotina are integrated structure made of two major kinds of tissues. The outer part, also called envelope or peridium, is exclusively made of maternal tissue whereas the inner part, the zygotic tissue is issued from fusion of the two parental haploid gametes [37]. To check whether the Δ*PaRid* sterility resulted from a peridium defect or from a developmental defect of the zygotic tissue, we set up trikaryon crosses involving the Δ*mat* strain [38,39]. Because the Δ*mat* strain lacks the genes required for fertilization, it does not participate either as male or female in sexual reproduction. However, the Δ*mat* mycelium is able to provide maternal hyphae required to build fruiting bodies. Consequently, the Δ*mat* strain can only complement mutants defective for the formation of the envelope but cannot complement zygotic tissue dysfunction. We observed that the Δ*mat* ; *mat+* Δ*PaRid*; *mat-* Δ*PaRid* trikaryons yielded micro-perithecia only (S6C Fig), equivalent in size and shape to the micro-perithecia generated by the *mat*+ Δ*PaRid* ; *mat*- Δ*PaRid* dikaryons. These results indicated a defect of zygotic tissues in Δ*PaRid* mutants. Furthermore, grafting Δ*PaRid* micro-perithecia onto a wild-type mycelium did not result in further development of the fruiting bodies and did not restore ascus production either (S4 Table), which disproves the hypothesis of a metabolic deficiency or a nutrient shortage being the cause of the observed Δ*PaRid* mutant strain defects [40]. Conversely, early stage (not yet mature) wild-type perithecia grafted onto Δ*PaRid* mycelia, continued to develop into fully mature perithecia and expelled the usual amount of ascospores confirming that the Δ*PaRid* mutant mycelium is able to provide all required nutrients.

### In the absence of PaRid in female gametes, sexual development is blocked before the dikaryon formation

As mentioned above, in *P. anserina*, fertilization results in formation of a plurinucleate ascogonium located inside the fruiting bodies. This ascogonium gives rise to dikaryotic cells that differentiate specialized cells (croziers) in which karyogamy leads to the formation of zygotes (S3 Fig). The diploid cells immediately enter meiosis to yield ascospores. In the homozygous Δ*PaRid* crosses, dissection of micro-perithecium contents (N=5) performed 48 hours post-fertilization showed plurinucleate ascogonial cells, but an absence of dikaryotic cells. This observation indicated that the sexual development was blocked before the formation of the dikaryotic cells (Fig 2D). Control experiment performed in the same conditions on perithecium contents collected from homozygous wild-type crosses (N=5), typically showed 20 to 30 dikaryotic cells per perithecium (Fig 2D). As mentioned above, heterozygous crosses where the Δ*PaRid* null allele was present in the female gamete genome and the wild-type *PaRid* allele in the male gamete genome resulted in the formation of micro-perithecia. When their content was dissected (N=5), no dikaryotic cells were observed as in the homozygous Δ*PaRid* crosses. By contrast, heterozygous crosses where the wild-type *PaRid* allele was present in the female gamete genome and the Δ*PaRid* null allele in the male gamete genome resulted in the formation of fully mature perithecia that contain dikaryotic cells (N=5). Altogether, these results show that a wild-type *PaRid* allele must be present in the haploid genome of *P. anserina’s* female gametes but not in the haploid genome of *P. anserina’s* male gametes for the dikaryotic cells to form.

Nevertheless, when homozygous Δ*PaRid* crosses are incubated from three to four weeks in the culture room (as opposed to the 96 hours post-fertilization needed for wild-type crosses to yield offspring), few asci were produced. A total of three asci were collected from 20 independent homozygous Δ*PaRid* crosses showing micro-perithecia only, whereas tens of thousands can typically be recovered from a single wild-type cross. Each of these 12 Δ*PaRid* dikaryotic ascospores displayed wild-type shape, germinated efficiently and generated *bona fide* Δ*PaRid* mutant strains.

### Complementation of the Δ*PaRid* mutants with ectopic alleles

To verify that the *PaRid* deletion was responsible for the sexual development arrest when absent from the female gamete haploid genome, we transformed a Δ*PaRid* strain with a *PaRid-HA* allele (see Material and Method section). Among the phleomycin-resistant transformants that were recovered (N=109), 84% showed a complete restoration of fertility (Table 1). Moreover, when the expression of *PaRid* was driven by the highly and constitutively active *AS4* promoter (*AS4-PaRid-HA* allele) [41], 67% of the phleomycin-resistant transformants (N=78) completed sexual development and produced ascospores (Table 1). Among these complemented transformants, no noticeable additional vegetative or reproductive phenotypes were observed (S4 Fig).

**Table 1.** Complementation experiments.

| Alleles | Fertile | Sterile |
| --- | --- | --- |
| *PaRid-HA* | 92 | 17 |
| *PaRid-GFP-HA* | 20 | 43 |
| *AS4-PaRid-HA* | 52 | 26 |
| *AS4-PaRid-GFP-HA* | 61 | 34 |
| *PaRid*<sup>C403S</sup>-*HA* | 0 | 89 |
| *AS4-PaRid*<sup>C403S</sup>-*HA* | 0 | 55 |
| *NcRID-HA* | 0 | 120 |
| *dmtA-HA* | 0 | 112 |
| *Masc1-HA* | 0 | 86 |

A collection of alleles was introduced in a Δ*PaRid* strain and subsequent transformants where crossed to a Δ*PaRid* strain of compatible mating type. Number of fertile versus sterile transformants is indicated in the table. When none of the transformants showed a restored fertility, PCR amplifications were performed to check the presence of full-length ectopic alleles. Those from which we cannot amplify the corresponding fragment were discarded. For each transformation experiments, expression of the HA-tagged proteins was assayed by western blot on a subset of transformants (see S7 Fig). See results section for details on allele features.

To investigate whether the putative enzymatic function of the PaRid protein was essential to *P. anserina*’s sexual development, we constructed the *PaRid*^C403S^*-HA* and the *AS4-PaRid*^*C403S*^*-HA* point-mutated alleles, which both encode a catalytically-dead PaRid protein [1]. To perform the methylation transfer, this cysteine residue of the conserved PCQ triad (proline-cysteine-glutamine) located in motif IV forms a covalent bond with the cytosines that will be modified (Fig 1D). This cysteine residue is invariant in all eukaryotic C5-cytosine methyltransferases and its substitution results in loss of activity in mammalian DNA methyltransferases Dnmt3A and Dnmt3B [42]. After transformation of a knockout Δ*PaRid* strain, independent phleomycin-resistant transformants were recovered (N=89 for the *PaRid*^C403S^*-HA* allele and N=55 for the *AS4-PaRid*^*C403S*^*-HA* allele). Although they presented at least one full-length copy of either the *PaRid*^C403S^*-HA* allele or the *AS4-PaRid*^*C403S*^*-HA*, none of these transformants showed any complementation of the Δ*PaRid* sterility (Table 1). For a subset of them, we confirmed that the corresponding point-mutated protein PaRid^C403S^ was readily expressed (S7 Fig). Again, these transformants could not be distinguished from Δ*PaRid* mutants when crossed to either wild-type strains or Δ*PaRid* mutant strains.

Finally, since the Masc1/RID proteins are phylogenetically and structurally related and they might play a conserved role during sexual development, at least for the *A. nidulans* and *A. immersus* orthologs, we transformed the *DmtA-HA* and *Masc1-HA* alleles but also the *NcRid-HA*, independently, under the control of a functional promoter, into the Δ*PaRid* background. None of the independent phleomycin-resistant transformants that were recovered showed restoration of sexual reproduction (N=120 for the *NcRid-HA* allele, N=112 for the *DmtA-HA* allele and N=86 for the *Masc1-HA* allele, Table 1), indicating that inter-species complementation is ineffective.

### PaRid cellular localization

To investigate the subcellular localization of PaRid, we expressed two GFP-tagged chimeric versions of this protein (Table 1 and Fig 3). The *PaRid-GFP-HA* allele was driven by its native promoter, while the *AS4-PaRid-GFP-HA* allele was driven by the highly and constitutively active *AS4* promoter. After transformation of a Δ*PaRid* strain, independent phleomycin-resistant transformants were recovered (N=63 for the *PaRid-GFP-HA* allele and N=95 for the *AS4-PaRid-GFP-HA* allele). Since 32% of the phleomycin-resistant strains obtained after transformation with the *PaRid-GFP-HA* allele and 64% of those obtained after transformation with the *AS4-PaRid-GFP-HA* allele showed a complete restoration of the Δ*PaRid* fertility defect (Table 1), both tagged-alleles were proven to be expressed and to encode functional proteins. Fluorescence under the native promoter was too weak to be monitored so that overexpression constructs were subsequently analyzed. All the complemented *AS4-PaRid-GFP-HA* transformants had a wild-type phenotype (*i.e.*, displayed no spurious phenotypes that could be linked to PaRid constitutive overexpression). The PaRid-GFP fluorescence was observed in the female gametes (ascogonia and protoperithecia, Fig 3A and B) but neither in mycelium nor in the male gametes (spermatia). This expression pattern is in line with both the Δ*PaRid* maternal sterility and the absence of *PaRid* ESTs in vegetative mycelium. Surprisingly, fluorescence could no longer be observed in croziers (formed around 30 hours post-fertilization) (Fig 3C), but signal resumed during ascospore formation (from 42 to 96 hours post-fertilization) (Fig 3D). In ascospores, PaRid-GFP was both nuclear and cytoplasmic.

**Fig 3.**
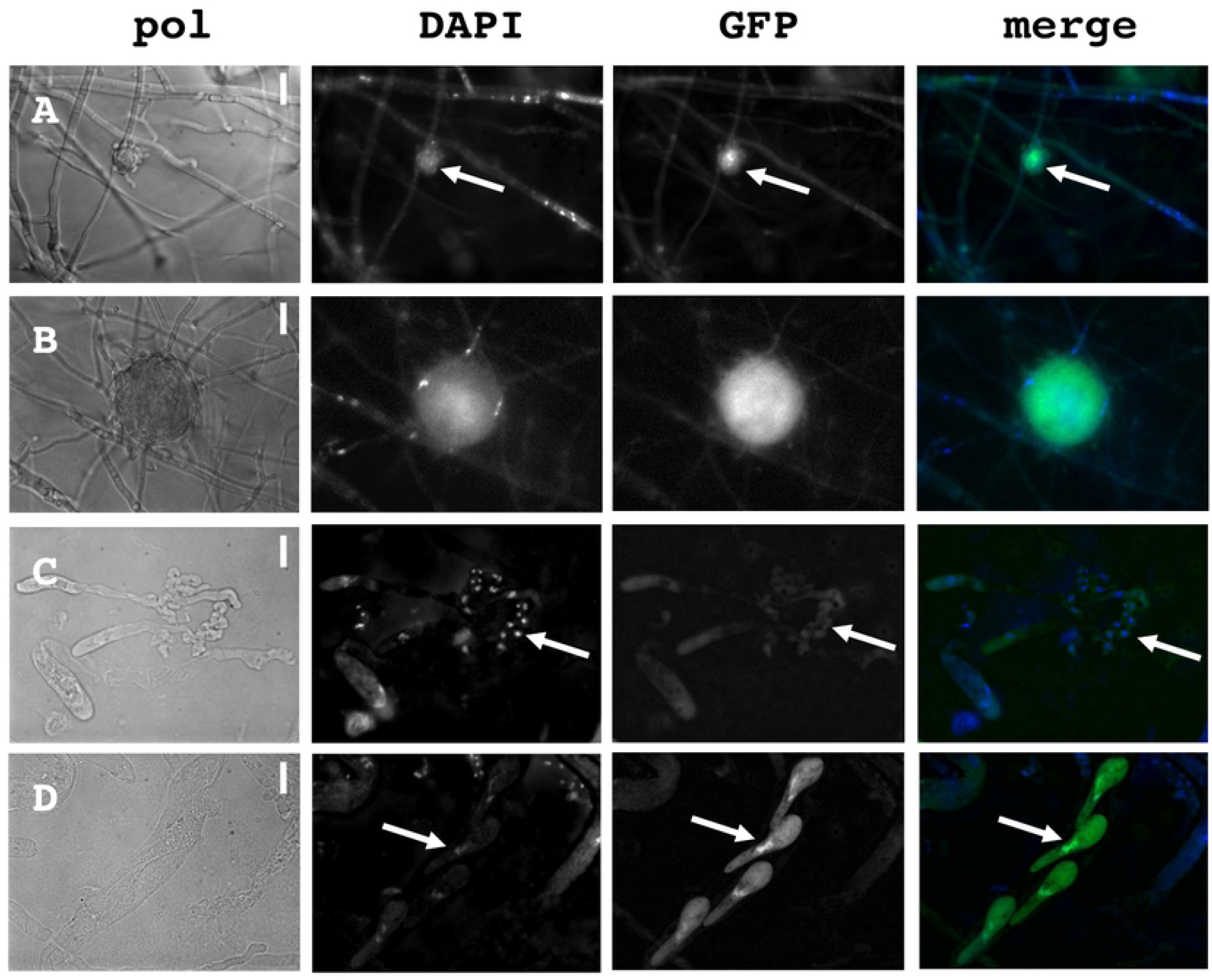
Expression and subcellular localization of PaRid during *P. anserina* life cycle. PaRid-GFP expression was assayed from a ΔPaRid:AS4-*PaRid*-GFP-HA strain showing wild type phenotypes. No GFP signal can be observed in mycelium, however, a significant and specific GFP signal can be found in ascogonia (A, white arrows are pointing at ascogonia) and protoperithecia (B). Surprisingly, no GFP signal is observed in the croziers (C, white arrows are pointing at croziers) but a strong signal can be noticed in the mature ascospores (D, white arrows are pointing at the nuclei of one ascospore), showing both a nuclear and cytoplasmic localization of PaRid. From left to right: bright-field (pol), DAPI staining, GFP channel and merge of the two latest. Scale bar: 1,5 µm (A and B), 5 µm (C and D).

Notably, such temporary loss of fluorescence in the *P. anserina’s* crozier cells was not an isolated observation concerning nuclear proteins [43]. One can hypothesize that this cell-specific drastic reduction of GFP brightness might result from either low pH or high oxiradical contents [44,45]. On the contrary, seeing some GFP signal further supports the detection of *PaRid* transcripts by RT-PCR up to 96 hours post-fertilization (S1A and B Figs), whereas in the microarray experiments, the ratio of ascogenous tissues versus vegetative tissues (perithecium envelope) might be low, thereby masking *PaRid*’s expression profile.

### Transcriptomic reprograming in response to Δ*PaRid* developmental arrest

We explored the transcriptional profile of the Δ*PaRid* micro-perithecia using *P. anserina’s* microarrays, representing 10556 predicted coding sequences (CDS) [46]. To this end, Δ*PaRid mat*+ female gametes were fertilized by wild-type *mat*-male gametes and micro-perithecia were collected 42 hours post-fertilization (T42). This time point was chosen to unravel the broadest set of differentially expressed genes, given that 1) micro-perithecia originating from Δ*PaRid* crosses do not form croziers and thus might stop to develop around 30 hours (T30) post-fertilization, 2) in wild-type crosses, croziers are formed from 30 hours (T30) to 42 hours post-fertilization (T42) afterward karyogamy can proceed (Fig 4A). To identify the differentially expressed CDS (DE CDS), we compared the transcriptional profile of the Δ*PaRid* micro-perithecia to that of the wild-type perithecia at both 24 (T24) and 30 (T30) hours post-fertilization (Bidard and Berteaux-Lecellier, GEO accession no. GSE104632), hypothesizing that these two time points likely correspond to the window of time that immediately precedes and/or spans the Δ*PaRid* developmental arrest. Doing so, we identified 451 CDS which expression was either down-regulated (217 CDS) or up-regulated (234 CDS) by a fold change (FC) ≥ 2 (Table S5) at both T24 and T30 time points. This set of DE CDS represented 4.4 % of *P. anserina’s* predicted CDS. Actually, none of the CDS had opposite patterns of differential expression at T24 and T30.

**Fig 4.**
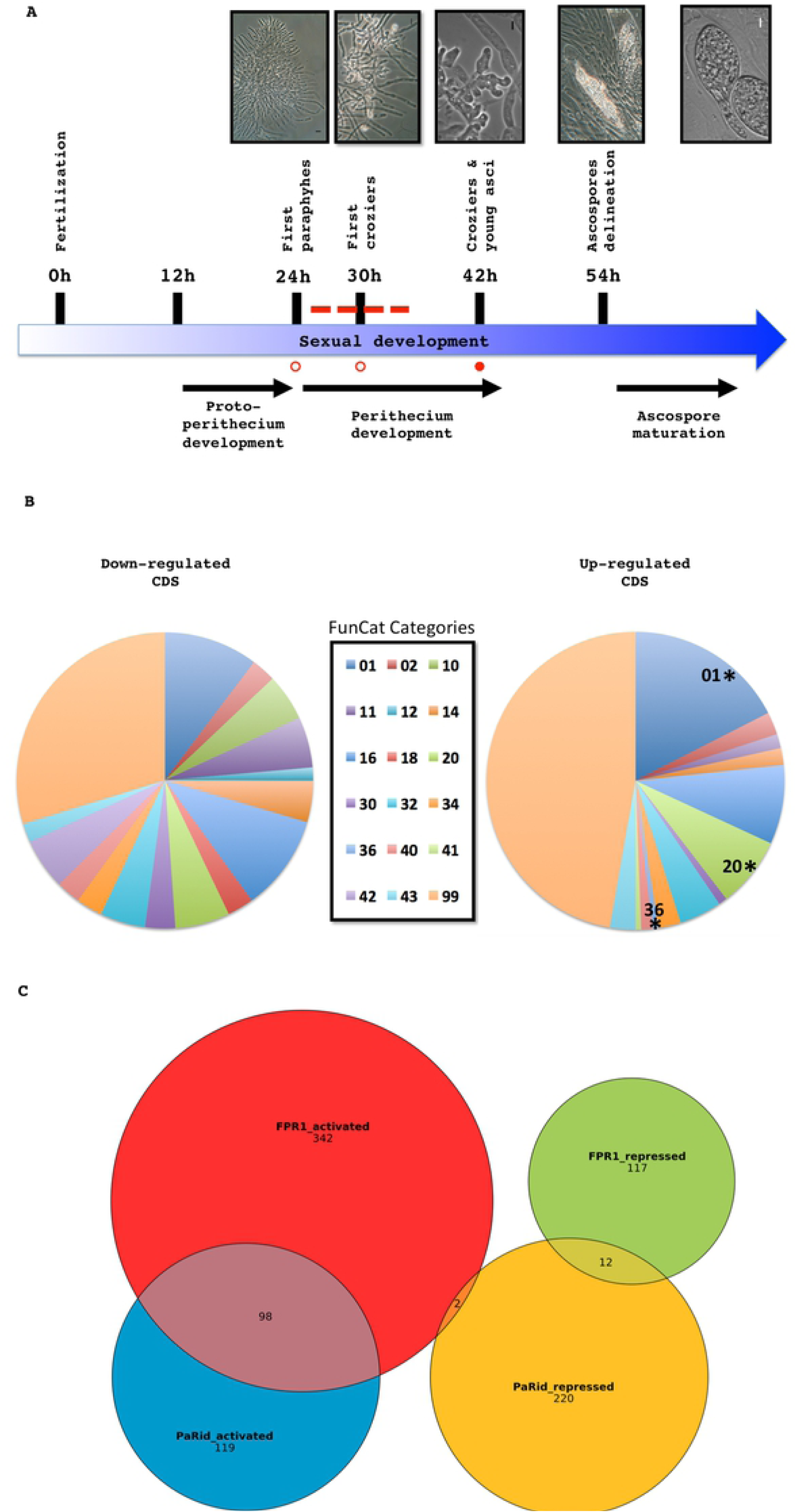
Transcriptomic analysis experimental design and data. (**A**) Schematic developmental time course from fertilization to ascospore maturation. Total RNA was extracted from wild-type perithecia at T24 and T30 (open circles) and from Δ*PaRid* micro-perithecia at T42 (solid circle); Dotted line: time frame during which the Δ*PaRid* developmental blockage might occur. Light microphotographs of the upper panel illustrate the various developmental steps indicated along the time course. **(B)** Functional categories in the down- and up-regulated CDS sets. Legend of pie charts corresponds to the FunCat categories, see Table 2 for details. Stars mark significantly enriched functional categories (p-value < 0.05). **(C)** Venn diagram of PaRid and FPR1 targets.

### Functional annotation of up- and down-regulated genes

CDS with no predicted function or domain were not over-represented in neither down-regulated (72 out of 217) nor up-regulated (80 out of 234) sets of DE CDS when compared to the rest of the genome (Table S5). The fraction of DE CDS showing *N. crassa* or *S. cerevisiae* orthologs was 69.6% and 32.2% respectively in the down-regulated set and 50.9% and 11.9% respectively in the up-regulated set. These observations show that DE CDS having an ortholog either in *N. crassa* or in *S. cerevisiae* were significantly less abundant in the up-regulated set than in the down-regulated set (p-value = 0.0455 and 3.10*10^-5^, respectively). This bias existed also when the up-regulated CDS having an ortholog either in *N. crassa* or in *S. cerevisiae* where compared to those of the complete *P. anserina’s* set of CDS (p-value = 0.0317 and 9.38*10^-8^, respectively). This might indicate that the down-regulated set belongs preferentially to the conserved fungal genome core whereas the up-regulated set appears more divergent and species specific. We then performed a FunCat analysis [47] to better characterize the function of the DE CDS (Fig 4B, Table 2). Approximately two-third of them belonged to the “Unclassified” category (category number 99) either in the up- or down-regulated sets. Among the 72 classified CDS of the down-regulated set, no FunCat categories were enriched (See Table 2). By contrast, the up-regulated set was enriched in CDS that fell in the “Metabolism” (category number 01) and “Cellular transport” categories (category number 20).

**Table 2.**
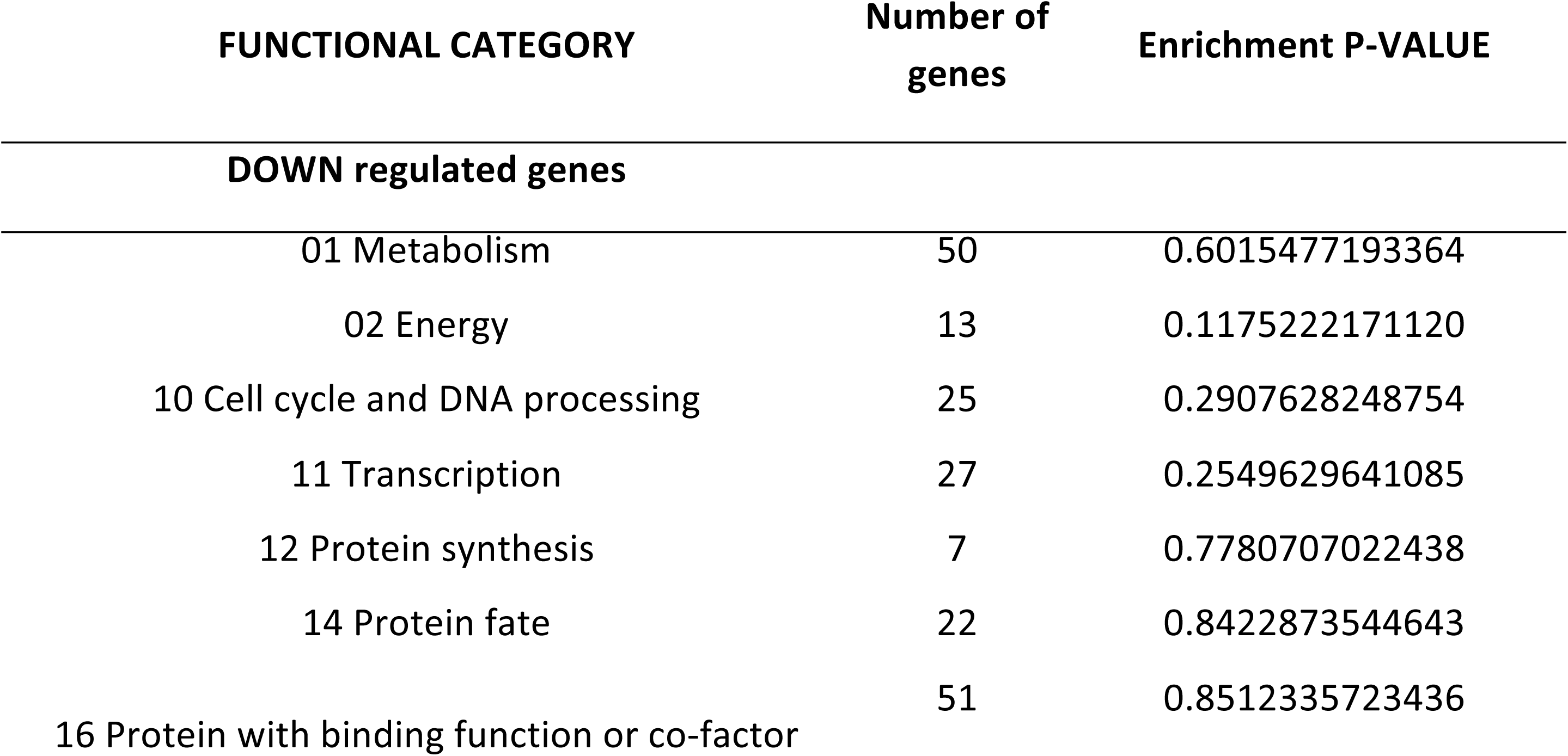
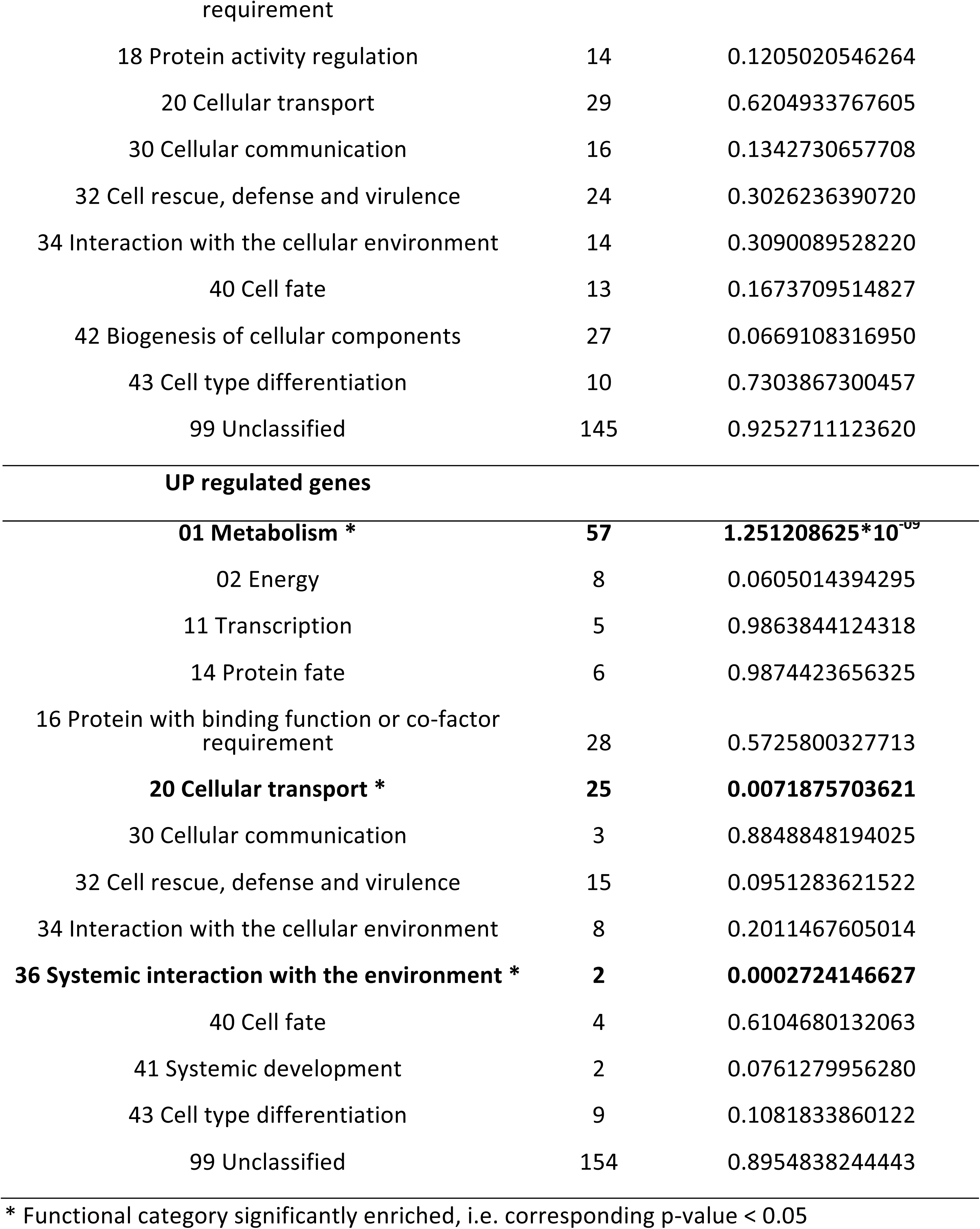
Functional category analysis (FunCat).

### Comparison of PaRid regulated genes and mating-type target genes shows a significant overlap

We compared the Δ*PaRid* DE CDS to the mating-type target genes identified previously [48]. *P. anserina* has a typical heterothallic mating-type structure with two idiomorphs. The *MAT1-1* idiomorph (*mat*-) is characterized by the *MAT1-1-1* gene, which encodes the FMR1 MATα-HMG domain containing protein, while the *MAT1-2* idiomorph (*mat*+) is composed of the *MAT1-2-1* gene, which encodes the FPR1 MATA_HMG domain containing protein (reviewed in [49]). Both proteins are transcription factors essential for fertilization in heterothallic Pezizomycotina and development of the fruiting bodies (reviewed in [37]). Microarray comparisons of wild-type *mat+* versus *fpr1*^*-*^ mutant strains, and wild-type *mat-* versus *fmr1*^*-*^ mutant strains revealed 571 and 232 target genes of FPR1 and FMR1, respectively [48]. The authors have determined that among the FPR1 target genes, 442 are activated and 129 are repressed. Similarly, among the FMR1 target genes, 151 are activated and 81 are repressed. Comparing these activated and repressed mating-type target genes with the genes down- and up-regulated in the Δ*PaRid* mutant strain showed a significant overlap (Fig 4C, S5 Table). The 217 Δ*PaRid* down-regulated CDS contained 98 FPR1 activated targets, which was clearly indicative of a strong enrichment (p-value = 5.40*10^-46^). FPR1 acts also as a repressor: accordingly we found 14 FPR1 repressed targets among the 234 Δ*PaRid* up-regulated CDS, which did not correspond to any enrichment (p-value > 0.05) (Fig 4C, S5 Table). Strikingly, we noticed that there is no CDS that would be both down-regulated in the Δ*PaRid* background and repressed by FPR1, while only two CDS were up-regulated in the Δ*PaRid* background and activated by FPR1 (Fig 4C, S5 Table). This observation indicates a strong congruence of the regulatory pathways of PaRid and mating-type gene FPR1. Conversely, only 17 FMR1 targets were identified in the 541 PaRid regulated CDS. This low number of FMR1 targets (*mat-* idiomorph) is consistent with the Δ*PaRid mat+* mutant strain being used as the female partner in the microarray experiments (Table 3) and our above observation that PaRid is dispensable in the male partner (here *mat*-). Out of the 17 FMR1 targets, five were found in the Δ*PaRid* down-regulated CDS set whereas 12 were found in the Δ*PaRid* up-regulated CDS set (S5 Table).

**Table 3.** Crosses used in transcriptome analysis.

| Denomination | Crosses | Female strain | Male strain | Sampling time |
| --- | --- | --- | --- | --- |
| **Wild type cross** | *S mat+* x *S mat-* | *S mat+* | *S mat-* | T24 and T30 |
| **$\Delta PaRid$ cross** | *S mat+* $\Delta PaRid$ x *S mat-* | *S mat+* $\Delta PaRid$ | *S mat-* | T42 |

### Most of the down-regulated DE CDS are involved in developmental pathways

The down-regulated Δ*PaRid* CDS set was enriched in transcription factors (TFs, p-value = 1.02*10^-4^). Notably, out of the 17 TFs identified (S6 Table, S8 Fig), 11 were also FPR1 targets. Furthermore, Pa_1_16860 and Pa_3_1720 orthologs in *N. crassa* [50,51] and *A. nidulans* [52,53] influence sexual development (S6 Table). On the same note, the *N. crassa* ortholog of Pa_1_22930 plays a significant role in sexual development [51]. Yet, we did not identify in this down-regulated CDS set, the *P. anserina’s* orthologs of most of the *N. crassa* TFs that have been found to be differentially expressed during sexual development [50,54–57].

We also noticed three enzymes related to NAD/NADP oxidoreduction and belonging to the developmental class: a NADPH dehydrogenase (Pa_6_6330), a NADP-dependent oxidoreductase (Pa_5_11750) and a glycerol-3-phosphate dehydrogenase [NAD+] (Pa_1_6190), as well as proteins involved in cellular signal transduction by regulating the phosphorylation status of the intracellular inositol trisphosphate messenger, including PaIPK2 [58] (Pa_5_1490, Pa_6_9890, Pa_1_18990). Interestingly, inositol phosphates are required for both fruiting body number and proper development in *Sordariales* [59].

In this down-regulated set, we also identified widely conserved CDS regulating cell division: Ras GTPase-activating proteins (Pa_1_10960 and Pa_6_7140), Rho-GTPase-activating protein (Pa_7_10800), cell division control protein (Pa_3_3430), replicating licensing factor (Pa_4_8520) and serine/threonine-protein kinases (Pa_1_9100 and Pa_7_9140). In line with active DNA replication, we also spotted some enzymes or cofactors involved in the nucleotide metabolism, such as a 3’,5’-cyclic-nucleotide phosphodiesterase (Pa_7_2860), a dimethyladenosine transferase (Pa_2_13220) and two guanine nucleotide-binding protein alpha-2 subunits (Pa_2_10260, Pa_5_11490). The reduced expression of genes involved in promoting cell division was clearly in line with the growth arrest of the Δ*PaRid* micro-perithecia. The developmental category may be more informative to uncover some CDS acting downstream of the PaRid network. In this category, we identified the PEX3 peroxisomal biogenesis factor (Pa_7_8080). The crucial role of β-oxidation and the glyoxylate cycle during sexual development has already been documented [60]. We also found a VelvetA-like-1 protein (VeA, Pa_3_6550) [61]. Present in many fungi, the Velvet protein complex seems to have expanded its conserved role in developmental programs to more specific roles related to each organism’s needs [62,63]. Likewise, the *IDC1* gene that was found down-regulated in this study is required for both cell fusion and development of the envelope of the fruiting bodies. The list of the down-regulated CDS includes a putative protein presenting a fascilin (FAS1) domain. Such extracellular domain is thought to function as a cell adhesion domain [64] and therefore might play a key role in the development of multicellular structures. A further connection to cell shape dynamics and cytoskeleton was found with the down-regulation of an annexin protein (Pa_6_1130). This calcium-dependent phospholipid-binding protein family has been linked with membrane scaffolding, organization and trafficking of vesicles, endocytosis, exocytosis, signal transduction, DNA replication, etc. The *N. crassa* homolog of this annexin (NCU04421) was shown to be up-regulated during asexual sporulation [65].

The gene expression regulation class mostly contained general regulating factors as CDS encoding chromatin remodeling proteins. As down-regulated chromatin remodeling factors, we identified two histone-lysine N-methyltransferases, Pa_7_3820 homologous to Set-9/KMT5, which methylates the lysine 20 of histone H4 [66] and Pa_3_3820 homologous to set-4, one acetyltransferase (Pa_3_10520), one ATP-dependent RNA helicase (Pa_4_8200) and one SNF2 family ATP-dependent chromatin-remodeling factor (Pa_7_9570). Some CDS encoding DNA/RNA processing factors were also found down-regulated in the Δ*PaRid* mutant micro-perithecia. As such, we identified a putative cruciform DNA recognition protein (Pa_2_440), an ATPase involved in DNA repair (Pa_6_4260), a SIK1-like RNA-binding protein (Pa_5_12950) and the telomere length regulator protein rif1 (Pa_1_3890). In mammals as in yeast, Rif1 is required for checkpoint-mediated cell cycle arrest in response to DNA damage [60].

### Analysis of the up-regulated CDS set uncovers enrichment in the “Metabolism” and “Cellular transport” functional categories

Half of the 26 most up-regulated CDS were putative proteins of unknown function devoid of conserved domains (Table S5), included the one showing the highest FC (Pa_4_5390). Two CDS encoded enzymes involved in the metabolism of amino acids (Pa_4_140 and Pa_1_5740), two were encoding transporters (Pa_3_420 and Pa_6_11600) and three CDS were involved in secondary metabolism (Pa_4_4580, Pa_5_11000 and Pa_5_720, see below).

More importantly, CDS belonging to primary metabolism were enriched in this up-regulated set (p-value = 2.35*10^-9^). For instance among the CAZymes, 11 glycoside hydrolases (GH) were found in this set (enrichment p-value = 0.0186). The predicted secondary metabolite clusters identified in this study are mostly composed of CDS encoding proteins of unknown function. Nonetheless, some of them contain CDS encoding putative secondary metabolism related functions: HC-toxin synthetases (Pa_3_11193, Pa_3_11220), cytochrome P450 proteins (Pa_3_2900, Pa_4_4580, Pa_4_4570, Pa_1_9520, Pa_6_7810), polyketide synthase (Pa_5_11000), multidrug efflux systems (Pa_3_11220, Pa_4_3775), trichodiene oxygenase (Pa_3_5540) and an O-methylsterigmatocystin oxidoreductase (Pa_2_7080). If they are not the result of cellular stresses induced in the Δ*PaRid* mutants, these secondary metabolites could act as secondary messengers during *P. anserina* sexual development. CDS encoding transporters were also over-represented in the up-regulated CDS set (p-value = 0.0071, see Table 2), as if arrest of perithecium development would generate cellular flux. Surprisingly, we also identified eight CDS encoding HET domain-containing proteins (Pa_2_4570, Pa_2_9350, Pa_2_8040, Pa_3_2610, Pa_5_1080, Pa_5_7650, Pa_6_1970, Pa_6_6730) (enrichment p-value = 0.0390). The HET gene family is known to prevent cell fusion in filamentous fungi by inducing cell death program when genetically different nuclei cohabit in a common cytoplasm [68]. Formation of heterokaryotic mycelia and potential incompatibility responses to non-self are vegetative processes that might be repressed through the PaRid network.

## Discussion

Sexual reproduction is considered to be essential for long-term persistence of eukaryotic species [69]. Only a few asexual lineages are known to persist over a long period of time without sex [70], most eukaryotes engaging sexual reproduction at some point in their life cycle. Studies have shown that sex reduces the accumulation of deleterious mutations compared to asexual reproduction [71] but also provide a faster adaptive response, by bringing together favorable gene combinations [72,73]. In multicellular eukaryotes, sexual reproduction is controlled by strict mechanisms governing which haploids can fuse (mating) and which developmental paths the resulting zygote will follow. These strict mechanisms are both genetically and epigenetically determined. Among the epigenetic modifications that control gene expression, DNA methylation reprogramming allows cells to shape their identity by launching and maintaining differential transcriptional programs in each cell type. In mammals, specific differential genomic DNA methylation patterns of parental gametes, known as ‘DNA methylation imprints’ are not essential to karyogamy but to zygotic development [74] while global genomic demethylation in *A. thaliana* results in male-fertile but female-sterile plants [23,24]. It has been hypothesized that the regulation of gene expression by DNA methylation during the development of higher eukaryotes may have been acquired from ancestral mechanisms of genome defense against invasive repetitive elements such as transposons. Some of the multicellular fungi are endowed with a homology-based genome defense system (RIP or RIP-like) exhibiting epigenetic features in addition to a functional link to sexual development [11,18]. Nonetheless, very little is known about epigenetic regulation of gene expression and development in multicellular fungi, although animal and fungi are each other’s closest relatives. In this study, we took advantage of a model ascomycete that displays on-going RIP but no DNA methylation [14], to explore the role of PaRid, a DMT-like protein, part of the Masc1/RID family, conserved in fungi [11,12].

### PaRid is involved in the formation of croziers

We report that PaRid plays a central role in the mid-time course of sexual development of *P. anserina*. Homozygous Δ*PaRid* crosses, as heterozygous ones, can perform fertilization, which depends on the recognition of male cells by female organs. Therefore PaRid is not involved in the early steps of sexual development. However, the *PaRid* wild-type allele must be expressed in the female gametes for further perithecium development. The most conspicuous developmental step after fertilization is the formation of the dikaryotic cells, which is preceded by the division of parental nuclei in a syncytium (ascogonial plurinucleate cell). If PaRid is absent from the maternal lineage, the perithecia stop to grow prematurely (micro-perithecia phenotype) and no croziers emerge from the biparental plurinucleate ascogonial cells that they contain. Furthermore providing surrogate maternal tissue either by grafting the mutant micro-perithecia to a wild-type mycelium or by introducing a Δ*mat* partner to the Δ*PaRid mat+* ; Δ*PaRid mat-* dikaryon did not rescue the maturation defect of the fruiting bodies, as previously shown for some of the *P. anserina* female sterile mutants [39,75–77]. This result demonstrates that the Δ*PaRid* mutant sterility is not due to a perithecium envelope building failure, nor to an improper feeding of the maturing fruiting body, but to the lack of ascogenous tissue. To date, among the *P. anserina* sterile mutants that have been studied, these macroscopic (micro-perithecia) and microscopic (no dikaryotic cells) phenotypes are both found only in the Δ*Smr1* mutant [78]. SMR1, MAT1-1-2 in the standard nomenclature [79], is a conserved protein of unknown molecular function, which is required for the initial development of the dikaryotic stage of both *Gibberella zeae* [80] and *Sordaria macrospora* [81]. A noticeable difference between *PaRid* and *SMR1*/*MAT1-1-2* is that the former must be present in the maternal lineage (*mat+* or *mat-*), while the latter is present only in the *mat-* nucleus (male or female). Genetic observations support the idea that SMR1 diffuses from *mat-* to *mat+* nuclei inside the fruiting body, even if *mat-* nuclei come from a male gamete [82]. In contrast, our observations based on the tagging of PaRid with the GFP indicate that this gene is not expressed in the male gamete, and complementation experiments suggest that *PaRid* is not expressed in nuclei coming from male gametes inside the perithecia. It has been proposed that SMR1 releases the developmental arrest following inter-nuclear recognition between two compatible nuclei in the plurinucleate ascogonial cell and could consequently trigger crozier formation [83]. Besides, in contrast to the Δ*PaRid* and Δ*Smr1* deletions, the previously identified *P. anserina* mutants affecting the zygotic lineage (Smr2, Cro1 and Ami1), generate either uninucleate croziers [33,34,78] or plurinucleate croziers [84]. Consequently, because the corresponding mutants do indeed form these early hook-shaped structures, PaRid probably acts at earlier developmental stages than Smr2, Cro1 and Ami1, in line with its expression pattern. Taken together, these data suggest that SMR1 releases the developmental arrest following inter-nuclear recognition by diffusing from the mat- to mat+ nucleus, while *PaRid* is a maternal gene that is required for further development, especially the formation of the croziers.

### PaRid is an activator of the mid-time course of *P. anserina*’s sexual development

In an effort to uncover the PaRid regulatory network and identify potential co-factors, we performed a transcriptomic analysis of mutant crosses. Strikingly, this study established that almost half of the CDS activated by PaRid (45.16% of the down-regulated set) are also activated by the FPR1 transcription factor [48]. In agreement with the fertilization ability of Δ*PaRid*, the mat+ prepropheromone gene and the mat+ pheromone receptor gene are not included in the set of genes down-regulated in Δ*PaRid* background. Consequently, we concluded that 1) PaRid is a key actor of the middle steps of *P. anserina* sexual development; 2) PaRid along with FPR1 is involved in the developmental pathway that follows fertilization and leads to formation of dikaryotic cells; 3) PaRid, as FPR1, is an activator of this pathway. Furthermore, the expression of *FPR1* is not deregulated in the Δ*PaRid* mutant background, and conversely the expression of *PaRid* is not deregulated in the *fpr1*^*-*^ mutant background [48]. This result suggests that PaRid acts neither upstream nor downstream of FPR1, but rather as an independent branched pathway (Fig 5). The PaRid-regulated CDS set includes many FPR1 targets (n = 112) and few FMR1 targets (n = 17). This observation further supports the maternal effect of the *PaRid* deletion, as we have used a *mat+* (FPR1) strain as a maternal strain in our experiments. Determination of mating-type gene targets was performed at the stage of competence for fertilization [48], a stage belonging to the very early steps of sexual development of perithecia. As the mating-type gene targets are likely to vary according to the developmental stage [83], their determination in the middle steps of fruiting-body development would be more appropriate to the comparison with those of PaRid and might reveal even more extensive overlap. MAT1-2-1 (FPR1) and MAT1-1-1 (FMR1) are ubiquitous in Pezizomycotina (reviewed in [49]). We propose that the overlap of mating-type targets genes and PaRid-regulated genes is present in all fungi in which the inactivation of *Rid* results in sexual development defect. At present, *Rid* linked sexual defect were described in *A. immersus* [11] and *A. nidulans* [18]. Unfortunately, neither mating type target genes nor *Rid* regulated genes were available in these fungi to test our proposal.

**Fig 5.**
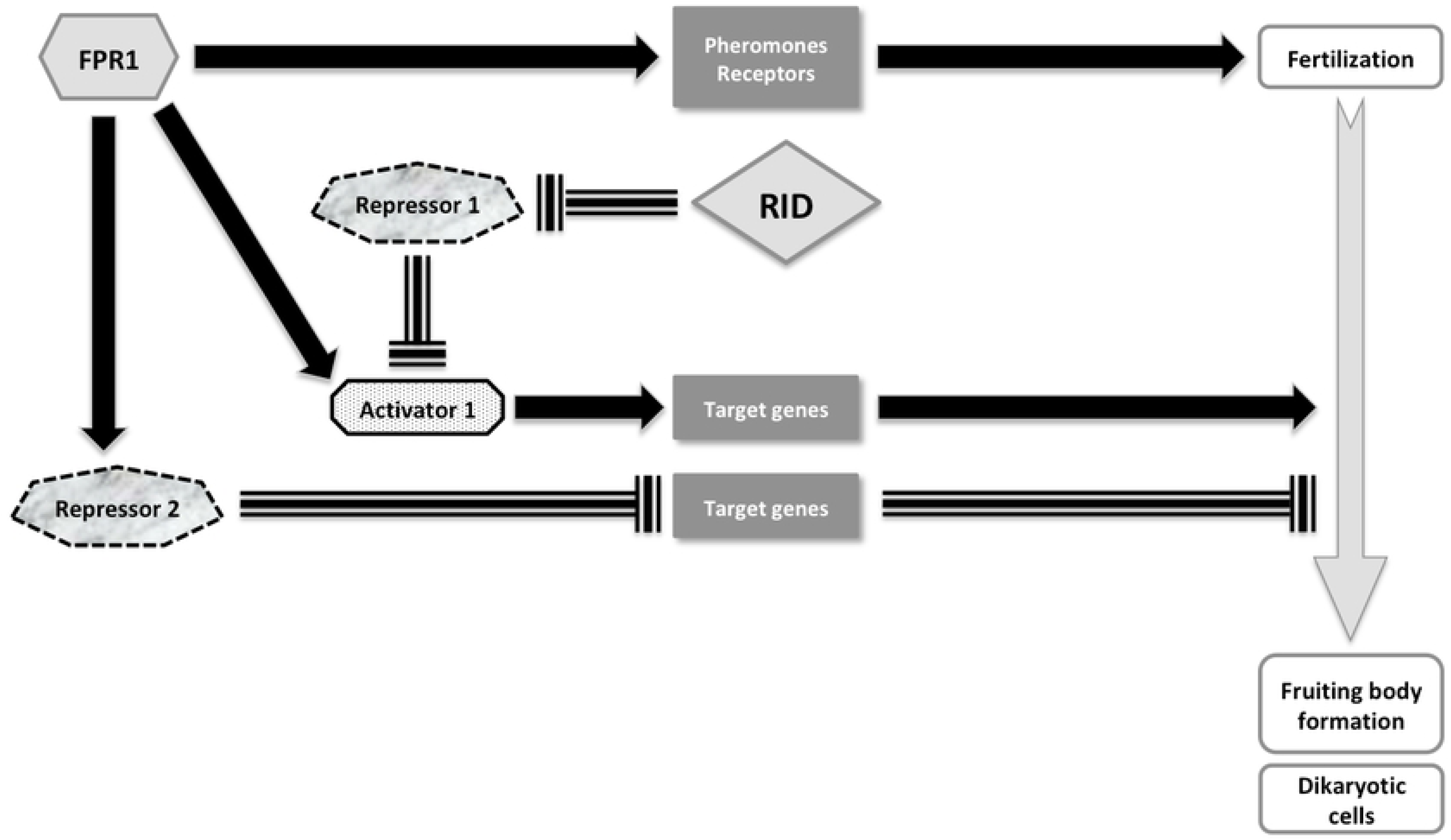
Schematic representation of PaRid & FPR1 developmental pathways during sexual development. FPR1, a MATα-HMG transcription factor is essential to fertilization and development of fruiting bodies. This mating type protein can act either as an activator or as a repressor. This study established that PaRid shares part of the FPR1 positive regulatory circuit, which is at work after fertilization to build the fructification and to form the dikaryotic cells. As the methyltransferase activity is required for PaRid function, we hypothesized that PaRid might repress a repressor. Solid black arrow is indicative of activation. Dashed T-line is indicative of repression. Solid grey arrow represents the sexual development time line.

However, this list of DE CDS did not draw a clear picture of a potential epigenetic regulation process, if any, in which PaRid could be involved, nor the nature of the Repressor 1 (Fig 5). Notably, neither the constitutive heterochromatin factors (the putative H3K9me3 methylase: Pa_6_990, the ortholog of the heterochromatin protein 1: Pa_4_7200, the components of the DCDC complex of *N. crassa* [85]: Pa_6_250, Pa_7_5460, Pa_2_10970, Pa_3_6830) nor the facultative heterochromatin factors (Pa_1_6940 encoding the putative H3K27 methylase and the associated subunits of the PRC2 complex: Pa_3_4080, Pa_2_11390) are differentially expressed. Genes involved in the STRIPAK complex [86] were not differentially expressed either. In consequences, the nature of Repressor 1 cannot be hypothesized yet (Fig 5). Nonetheless, this study identified a set of TFs that might be candidate to enter the specific *PaRid* genetic network, since eight out of 17 TFs of the Δ*PaRid* down-regulated CDS set are not FPR1 targets. Additional functional studies on a selected subset of specifically DE CDS identified in this study are required to explore the PaRid downstream pathway(s) which are essential to sexual development.

By contrast to the down-regulated set, the up-regulated one only contains a handful of developmentally relevant CDS. The functional annotation of the up-regulated set points toward CDS involved in physiological responses to the cellular stress following micro-perithecium developmental arrest.

### One toolkit, different outcomes

The Δ*PaRid* mutant phenotype resembles the ones observed in *A. immersus* [11] and in *A. nidulans* [18], where development of croziers and of the subsequent ascospores are never observed in *Masc1* mutants or *dmtA* mutants, respectively. Remarkably, sexual development of homozygous *T. reesei rid* mutant crosses deleted for the *rid* homolog (*TrRid*) allows the formation of dikaryotic cells but is blocked during karyogamy (Wan-Chen Li and Ting-Fang Wang, personal communication). On the contrary, neither barren perithecia nor fertility defect are observed in *N. crassa* crosses homozygous for the *rid* null allele [12], although this fungus shows heavy DNA methylation of repeats subjected to RIP. The function of *rid* orthologs has also been addressed in fungal species that reproduce asexually (*Aspergillus flavus* [87], *Cryphonectria parasitica* [88], *Metarhizium robertsii* [89]). Interestingly, in these species, in addition to a decrease of DNA methylation contents, the absence of DMT-like fungal specific Masc1/RID proteins results in a large palette of phenotypes including reduction of clonal dispersion (conidiation and sclerotial production), defects of mycelium morphology, decrease in secondary metabolite production and/or virulence toward plant hosts. In *Magnaporthe grisea*, a pathogenic fungus that can reproduce through both sexual and asexual reproduction, deletion of the *rid* ortholog also has a negative impact on conidiation and mycelium morphology but not on virulence [74].

Altogether, these observations are puzzling. It has led us to consider why such a gene, known to be involved in RIP-like genome defense systems and conserved both in terms of presence and sequence identity, would play an essential role during the sexual development in organisms showing RIP mutations at low level, and very little genomic DNA methylation, if any [14]. One can wonder if RIP, as a genome defense mechanism operating after fertilization but before karyogamy, is not by itself a checkpoint required for the development of dikaryotic hyphae in *P. anserina, A. nidulans* and *A. immersus*, and for karyogamy in *T. reesei*. In this hypothesis, in the absence of the ‘genomic quality control’ of the two haploid parental nuclei performed by RIP before every single karyogamy, the sexual development might stop. In this study, it was not possible to test whether PaRid is also required for RIP since homozygous Δ*PaRid* mutant crosses were barren, except for rare sporadic asci (only 3 recovered to date) and RIP frequency is low in *P. anserina* [33]. Nevertheless, all of the 12 ascospores that we recovered from these mutant crosses were indistinguishable from those of wild-type: their shape and pigmentation were standard, they germinated efficiently and subsequent genetic analyses showed that they did not result from uniparental lineages. Consequently, if RIP functions as a checkpoint, it does not prevent viability of progeny that circumvents the function of PaRid. In *N. crassa*, after karyogamy, genome integrity is further checked through the meiotic silencing of unpaired DNA process (MSUD) [91]. MSUD proceeds during meiosis to reversibly silence genes contained in DNA segments that are not paired with their homolog during meiotic homolog pairing. Because evolution often scrambles chromosomal micro-syntheny, the MSUD prevents interspecific crosses. If deletion of *PaRid* results in RIP deficient strains, then inactivation of genome defense system acting on haploid parental nuclei has clearly different outcomes, since it would promote repeat spreading.

### Ancestral dual roles for DMT-like fungal specific Masc1/RID?

Alternatively, PaRid might play a dual role, possibly relying on two independent enzymatic functions: one as the central effector of RIP and the other as a positive regulator of gene expression during the early steps of sexual development. Our results show that the PaRid function relies on its methyltransferase activity, and that this enzymatic activity has to be present in the female partner. Ascogonia, the female gametes, are small structures of specialized hyphae that differentiate within the mycelium [37]. As such, they represented an extremely low fraction of the so-called “vegetative” tissue when four-day-old mycelium was collected to extract total RNA. This might be why no *PaRid* mRNAs were detected at this stage while genetic data further supported by cytoplasmic observations indicate that PaRid is specifically present in the female gametes. Besides, since driving its expression from a constitutive promoter did not result in any unusual phenotypes, regulation of *PaRid* expression, both in terms of timing and cell-type specificity, might proceed through post-transcriptional and/or post-translational mechanisms. Although we did not detect any evidence of antisense *PaRid* transcript as described in *A. nidulans* [18], regulation of expression through non-coding RNA (ncRNA) is widely conserved in mammals, drosophila, plants and *S. pombe* [92,93]). Therefore, we cannot rule out that the *PaRid* pattern of expression is also regulated by ncRNA. Alternatively, the PaRid half-life might be tightly regulated by the ubiquitin proteasome system (UPS) [94–97]. Again, this hypothesis remains to be further explored.

Masc1/RID fungal-specific DMTs-like form a quite ancient group of proteins, whose initial DNA-methyltransferase function may have evolved differently in distinct lineages. The predicted structures of RID homologues, although conserved, are not identical [98]. In particular, the Masc1/DmtA/TrRid/PaRid proteins include a compact catalytic domain with a short C-terminal extension (at most 133 amino acids for *P. anserina*) when compared to that of *N. crassa* (260 amino acids). Mutant analyses revealed that the Masc1/DmtA/TrRid/PaRid proteins fulfill both sexual developmental functions and genome defense (either RIP is still active or traces of RIP are present as relics in the genomes), whereas the *N. crassa* RID protein plays a more specialized role, limited to its genome defense function. Taken these observations altogether, one can wonder whether the long C-terminal extension of *N. crassa* RID is responsible for its restricted function. However, neither the *Masc1* allele, nor the *DmtA* allele, restored the fertility defect of Δ*PaRid* mutants. This may indicate that despite a structural and functional conservation of the Masc1/DmtA/TrRid/PaRid group of proteins, some species-specific co-factors would be required for enzymatic catalysis. To date, none of them has been identified, although it has been suggested that the DIM-2 ortholog of *M. grisea*, MoDIM-2, mediates the MoRID *de novo* methylation [90,99].

Finally, since the Masc1/DmtA/TrRid/PaRid group appears to have a non-canonical motif VI structure when compared to all other prokaryotic and eukaryotic C5-cytosine methyltransferases [1], it is possible that this class of enzymes has acquired exclusive catalytic and/or substrate properties [32]. Although they do not share the features of Dnmt2, the mammalian tRNA cytosine methyltransferase, one can hypothesize that some of these fungal DMT-like enzymes could methylate RNA substrate(s). If true, this might explain why no clear nuclear localization signal was found in the PaRid protein and provide a consistent molecular basis to explain the maternal effect of the Δ*PaRid* mutation. However, to date, only N6-methyladenosine (m6A), the most prevalent modification of mRNA in eukaryotes, has been linked to developmental functions. For example, the budding yeast N6-methyladenosine IME4 controls the entry of diploid cells into meiosis [100] while lack of the *A. thaliana* ortholog MTA leads to embryonic lethality [101].

Further functional studies on the fungal DMT-like proteins should also help understand whether the critical PaRid-related developmental program is a conserved feature of Pezizomycotina, secondarily lost by *N. crassa* during fungal evolution.

## Materials and methods

### Strains and culture conditions

The strains used in this study derived from the « S » wild-type strain that was used for sequencing [35,102]. Standard culture conditions, media and genetic methods for *P. anserina* have been described [40,103] and the most recent protocols can be accessed at http://podospora.i2bc.paris-saclay.fr. Construction of the Δ*mus51::su8-1* strain lacking the mus-51 subunit of the complex involved in end-joining of broken DNA fragments was described previously [104]. In this strain, DNA integration mainly proceeds through homologous recombination. Mycelium growth is performed on M2 minimal medium. Ascospores do not germinate on M2 but on a specific G medium. The methods used for nucleic acid extraction and manipulation have been described [105,106]. Transformation of *P. anserina* protoplasts was carried out as described previously [107].

### Identification and deletion of the Podospora *PaRid* gene

The *PaRid* gene was identified by searching the complete genome of *P. anserina* with tblastn [108], using RID (NCU02034.7) [12] as query. One CDS Pa_1_19440 (accession number CDP24336) resembling this query with significant score was retrieved. To confirm gene annotation, *PaRid* transcripts were amplified by RT-PCR experiments performed on total RNA extracted from developing perithecia (female partner *mat+*/male partner *mat-*) at either 2 days (T48) or 4 days (T96) post-fertilization using primers 5s-PaRid/3s-PaRid (See S1 Table). Sequencing of the PCR products did not identify any intron in the *PaRid* ORF, thus confirming the annotation (see S1B Fig).

Deletions was performed on a Δ*mus51::su8-1* strain as described in [76] and verified by Southern blot as described in [109]. Because *PaRid* is strongly linked to the mating-type locus, the gene deletion has been performed in the two mating-type background to get sexually compatible Δ*PaRid* mutants.

### Allele construction

Two plasmids were mainly used in this study: the pAKS106 plasmid which derived from the pBCPhleo plasmid [110] and the pAKS120 plasmid which derived from pAKS106. pAKS106 contains an HA tag sequence followed by the *ribP2* terminator at the unique *Cla*I and *Apa*I sites [111]. The pAKS120 was generated by cloning the promoter of the *AS4* gene, which is highly and constitutively expressed throughout the life cycle [41], at the unique *Not*I site. Thus, recombinant C-terminus HA-tagged proteins can be produced from pAKS106 and pAKS120, the expression of the corresponding fusion gene being either under the control of its own promoter (pAKS106) or under the control of the constitutive highly expressed *AS4* promoter (pAKS120). Because our allele construction strategy required high-fidelity DNA synthesis, all PCR amplifications were performed using the Phusion High Fidelity DNA polymerase (Thermo Scientific), according to the manufacturer’s protocol and the resulting alleles were systematically sequenced to ensure that no mutation was introduced during PCR amplification (see S1 Table for primers sequences) and that, when relevant, the HA-tag or GFP sequences were in the appropriate reading frame.

### Constructing the *PaRid-HA* and *AS4-PaRid-HA* alleles and their GFP-tagged version

To complement the Δ*PaRid::hph* mutant strain, the wild-type *PaRid* allele was cloned into the pAKS106 plasmid. To this end, the wild-type *PaRid* allele was PCR amplified from the GA0AB186AD09 plasmid [35], using the primers PaRIDHAFXbaI and PaRIDHAREcoRI (see S1 Table). The 2914-bp amplicon, which corresponds to 653-bp promoter sequence followed by the complete *PaRid* CDS, minus the stop codon, was digested with the *Xba*I/*EcoR*I restriction enzymes and cloned into the pAKS106 plasmid, also hydrolyzed with the *Xba*I/*EcoR*I restriction enzymes to obtain the *PaRid-HA* allele. In order to put the *PaRid* allele under the control of a highly and constitutively expressed promoter, we then constructed the *AS4-PaRid-HA* allele. To do so, the wild-type *PaRid* allele was PCR amplified from the GA0AB186AD09 plasmid [35], using the primers PaRIDAS4XbaI and PaRIDHAREcoRI (see S1 Table). This pair of primers was designed to amplify the complete *PaRid* ORF, minus the stop codon. The resulting 2263-bp amplicon was digested with the *Xba*I/*EcoR*I restriction enzymes and cloned into the pAKS120 plasmid, also hydrolyzed with the *Xba*I/*EcoR*I restriction enzymes.

To tag the PaRid protein with the Green Fluorescent Protein (GFP), we PCR amplified the ORF of the e*GFP* minus the stop codon with primers RIDGFPFEcoRI and RIDGFPRClaI (see S1 Table), using the pEGFP-1 plasmid (Clontech, Mountain View, CA) as template. The resulting 713-bp fragment was digested with *EcoR*I and *Cla*I and cloned in frame with the *PaRid* allele either into the PaRid-HA plasmid or the AS4-PaRid-HA plasmid (see above), previously digested with the same restriction enzymes. This yielded the chimeric *PaRid-GFP-HA* and *AS4-PaRid-GFP-HA* alleles.

### Constructing the site-directed mutant *PaRid*^**C403S**^***-HA***

The PaRid^C403S^ allele was constructed by *in vitro* site-directed mutagenesis. The GA0AB186AD09 plasmid harboring the wild-type *PaRid* allele was amplified by inverse PCR, using the divergent overlapping primers RIDmut1 and RIDmut2. The RIDmut1 primer leads to G-to-C substitution at position 1208 of the *PaRid* sequence (see S1 Table, marked in bold italics), that matches the converse C-to-G substitution located in the RIDmut2 primer (see S1 Table, marked in bold italics). The PCR amplicons were digested with the *Dpn*I enzyme (Fermentas), self-ligated and then transformed into *E. coli* competent cells. The *PaRid* allele was sequenced from several plasmids extracted from independent chloramphenicol resistant *E. coli* colonies. One allele showing no other mutation but the directed T**G**T (cysteine) to T**C**T (serine) transvertion was selected as the PaRid^C403S^ allele. The *PaRid*^*C403S*^ allele was then PCR amplified using the primers PaRIDHAFXbaI and PaRIDHAREcoRI. The 2914-bp amplicon, which corresponds to 653-bp promoter sequence followed by the complete *PaRid* ORF, at the exception of the stop codon, were digested with the *Xba*I/*EcoR*I restriction enzymes and cloned into the pAKS106 plasmid, also hydrolysed with the *Xba*I/*EcoR*I restriction enzymes. This resulted in the *PaRid*^C403S^*-HA* allele. Then, to get the *PaRid* allele under the control of a highly and constitutively expressed promoter, we constructed the *AS4-PaRid*^*C403S*^*-HA* allele. To do so, the *PaRid*^*C403S*^ allele was PCR amplified from the pAKS106-*PaRid*^*C403S*^*-HA*, using the primers PaRIDAS4XbaI and PaRIDHAREcoRI (see S1 Table). This pair of primers was designed to amplify the complete *PaRid* ORF minus the stop codon. The resulting 2263-bp amplicon was digested with the *Xba*I/*EcoR*I restriction enzymes and cloned into the pAKS120 plasmid, also hydrolyzed with the *Xba*I/*EcoR*I restriction enzymes. This yielded the pAKS120-*AS4-PaRid*^*C403S*^*-HA* plasmid. All the PCR-amplified fragments were sequenced.

### Constructing the interspecific *NcRID-HA, dmtA-HA* and *Masc1-HA* alleles

In order to introduce the *N. crassa* wild-type *rid* allele [12], into the *P. anserina* Δ*PaRid::hph* mutant strain, we constructed the pAKS106-rid-HA plasmid. To this end, the *rid* gene was PCR amplified from wild-type *N. crassa* genomic DNA, using primers NcRIDHAFNotI and NcRIDHARBamHI (see S1 Table). The resulting 3420-bp amplicon, which corresponds to 861-bp of promoter sequence followed by the complete *rid* ORF, minus the stop codon, was digested with the *Not*I/*BamH*I restriction enzymes and cloned into the pAKS106 plasmid, also hydrolyzed with the *Not*I/*BamH*I restriction enzymes. This resulted in the *NcRid-HA* allele.

A different strategy was used to clone the *DmtA* gene from *A. nidulans* [18] and the *Masc1* gene from *A. immersus* [11]. Indeed, because these fungi were more distantly related to *P. anserina* than *N. crassa*, the wild-type CDSs were cloned into the pAKS120 plasmid and thus fused with the *P. anserina* AS4 promoter. To do so, the *DmtA* CDS was PCR amplified from a wild-type *A. nidulans* genomic DNA, using primers dmtAFXbaI and dmtARBamHI (see S1 Table). The resulting 1898-bp amplicon, which corresponds to the complete *DmtA* CDS, minus the stop codon, was digested with the *Xba*I/*BamH*I restriction enzymes and cloned into the pAKS120 plasmid, also hydrolyzed with the *Xba*I/*BamH*I restriction enzymes. This resulted into the *DmtA-HA* allele. Similarly, the *Masc1* allele was PCR amplified from a wild-type *A. immersus* genomic DNA, using primers AiRIDAS4XbaI and AiRIDHAREcoRI (see S1 Table). The resulting 1666-bp amplicon, which corresponds to the complete *Masc1* CDS, minus the stop codon, was digested with the *Xba*I/*EcoR*I restriction enzymes and cloned into the pAKS120 plasmid, also hydrolyzed with the *Xba*I/*EcoR*I restriction enzymes. This resulted in the *Masc1-HA* allele. All PCR-amplified fragments were sequenced.

### Cytology & microscopy analysis

Perithecia were harvested from 12 to 96 h after fertilization. Tissue fixation was performed as in [112]. Pictures were taken with a Leica DMIRE 2 microscope coupled with a 10-MHz Cool SNAP_HQ_ charge-coupled device camera (Roper Instruments), with a z-optical spacing of 0.5 mm. The GFP filter was the GFP-3035B from Semrock (Ex: 472nm/30, dichroïc: 495nm, Em: 520nm/35). The Metamorph software (Universal Imaging Corp.) was used to acquire z-series. Images were processed using the ImageJ software (NIH, Bethesda).

### Phenotypic analyses

Spermatium counting was performed as follows: each strain was grown on M2 medium at 27°C for 21 days. To collect spermatia, cultures were washed with 1.5 mL of 0.05% Tween 20 in sterile water. Numeration proceeded through Malassez counting chamber. Grafting was assayed as in [40]. Western blot analyses were performed on perithecia grown for two days, as in [111]. We used the anti-HA high affinity monoclonal antibody from rat to recognize the HA-peptide (clone 3F10, ref 11 867 423 001, Sigma-Aldrich).

### Phylogenetic analysis

Orthologous genes were identified using Fungipath [113] and MycoCosm portal [114] and manually verified by reciprocal Best Hits Blast analysis. Sequences were aligned using MUSCLE (http://www.ebi.ac.uk/Tools/msa/muscle/) and trimmed using Jalview to remove non-informative regions (i.e. poorly aligned regions and/or gaps containing regions). Trees were constructed with PhyML 3.0 software with default parameters and 100 bootstrapped data set [115]. The tree was visualized with the iTOL version 4.3 (http://itol.embl.de/). Functional annotation was performed using InterPro 71.0 (http://www.ebi.ac.uk/interpro/search/sequence-search), Panther Classification System version 14.0 (http://www.pantherdb.org/panther/), PFAM 32.0 (http://pfam.xfam.org/), Prosite (http://prosite.expasy.org/). Proteins were drawn using IBS v1.0.2 software (http://ibs.biocuckoo.org/)

### RNA preparation for microarray

The male partner was grown for 6 days at 27°C on M2 medium. Then, spermatia were collected by washing the resulting mycelium with 1.5 mL of H_2_O per Petri dish (10^4^ spermatia / ml). The female partner strains were grown for 4 days at 27°C on M2 covered with cheesecloth (Sefar Nitex 03-48/31 Heiden). The sexual development time-course experiment started when female partners were fertilized by spermatia (1.5*10^4^ spermatia / cross). This time point was referred to as T0 (0 hour). By scraping independent crosses, samples of 20 to 100 mg of growing perithecia were harvested 24 hours (T24) and 30 hours (T30) after fertilization from wild-type crosses and 42 hours (T42) after fertilization from Δ*PaRid* crosses and immediately flash-frozen in liquid nitrogen. For each time point, we collected three biological replicates originating from three independent crosses. The frozen samples materials were grinded using a Mikro-dismembrator (Sartorius, Goettingen, Germany). Total *P. anserina* RNA were extracted using the RNeasy Plant Mini Kit (Qiagen, Hilden, Germany) with DNase treatment. The quality and quantity of total RNA was assessed using a Nanodrop spectrophotometer (Nanodrop Technologies, Wilmington, USA) and the Bioanalyzer 2100 system (Agilent Technologies, Santa Clara, USA). This protocol was performed on three genetically distinct crosses as specified in Table 3.

### Labelling and microarray hybridization

Gene expression microarrays for *P. anserina* consisted of a custom 4×44 K platform (AMADID 018343, Agilent, Santa Clara, USA) containing 10,556 probes on each array with each probe in four replicates. Microarray hybridization experiments including target preparation, hybridization and washing were performed as described in [46]. The experimental samples were labeled with the Cy5 dye and reference sample with the Cy3 dye. The reference sample is a mixture of RNA extracted from different growth conditions.

### Microarray data acquisition, processing and analyses

Microarrays were scanned using the Agilent Array Scanner at 5µm/pixel resolution with the eXtended Dynamic Range (XDR) function. Array quality and flagging were performed as described [116]. Pre-processing and data normalization were performed with Feature Extraction (v9.5.3) software (Agilent technologies), with the GE2-v4_95_Feb07 default protocol. Statistical differential analysis were done using the MAnGO software [117], and a moderate *t*-test with adjustment of p-values [118] was computed to measure the significance with each expression difference. Differential analyses have been performed between the wild-type cross at 24 (T24) and 30 (T30) hours after fertilization and the mutant cross. Fold change values (FC) were calculated from the ratio between normalized intensities at each time of the time course. Differentially expressed (DE) coding sequences (CDS) in the Δ*PaRid* cross were defined as those whose adjusted *P*-values are inferior to 0,001 and the absolute value of FC higher than 2. The resulting list was called DE CDS specific of Δ*PaRid* cross. Microarray data reported in this paper have been deposited in the Gene Expression Omnibus database under the accession no. GSE104632. Functions of genes identified as up- or down-regulated in transcriptomics data were explored using FunCat functional categories (Level 1, as described in http://pedant.gsf.de/pedant3htmlview/html/help/methods/funcat.html). To assess whether any of the functions were observed in up- or down-regulated lists of genes at a frequency greater than that expected by chance, p-values were calculated using hypergeometric distribution as described in [119].

## Author Contributions

Conceived and designed the experiments: RD, FB, FM. Performed the experiments: HT, RB, FB, ABJ, PG, VB, FC, FM. Analyzed the data: PG, HT, RD, FB, FM. Contributed reagents/materials/analysis tools: PG, HT, RB, FB, ABJ, VB, FC, FM. Wrote the paper: FM.

### Acknowledgments

We acknowledge the technical assistance of Sylvie François. We are thankful to Philippe Silar for providing resources to start the project as well as for fruitful scientific discussion and to Ting-Fang Wang for sharing unpublished data and stimulating scientific discussion. We thank Nadia Ponts for performing the DNA methylation assays and Gaëlle Lelandais for performing statistics. We thank Cécile Fairhead, Tatiana Giraud, Marc-Henri Lebrun, Nadia Ponts and Annie Sainsard-Chanet for critical reading of the manuscript and fruitful discussions. P.G., F.C. and F.M. were supported by grants from UMR8621, UMR9198 and Agence Nationale de la Recherche Grant ANR-05-Blan-O385-02. We thank the Podospora Consortium for freely providing GA0AB186AD09 and of GA0AB186AD09.

## Supporting information

**S1 Fig. Pattern of expression of *PaRid*. (A)** Average expression profiles (y-axis) of *PaRid* (Pa_1_19440) during sexual development (x-axis, hours). **(B)** Amplification of *PaRid* transcripts (2296 bp) by RT-PCR. MW: GeneRuler DNA Ladder Mix (Thermo Fisher Scientific), RT T48, RT T96: RT-PCR performed on RNA extracted from 2 days or 4 days post fertilization developing perithecia, gDNA: genomic DNA, NRT: PCR performed on RNA extracted from 2 days or 4 days post fertilization developing perithecia, Neg: No RNA. See (Materials and methods section for details). **(C)** Coverage of RNA-seq mapped reads at the *PaRid* locus [36]. RNA-seq experiments were performed on RNA extracted from non-germinated ascospores (Ascospores), eight hours germinating ascospores (Germinated ascospores), 1-day- or 4-day-old mycelia, 2 days or 4 days post fertilization developing perithecia.

**S2 Fig. Molecular characterization of knock-out mutants by Southern-blot hybridization.** Schematic representations of the endogenous and disrupted loci are given in (S2A and S2B). Replacement by homologous recombination of the wild type *PaRid* allele by the disrupted Δ*PaRid* allele results in the substitution of a 3.2 kb *Pst*I fragment by a 2.3 kb *Pst*I fragment as revealed by hybridization of the 5’UTR digoxygenin-labeled probe (S2A) and in the substitution of a 2.8 kb *Pst*I fragment by a 4.0 kb *Pst*I fragment as revealed by hybridization of the 3’UTR digoxygenin-labeled probe (S2A).

**S3 Fig. Major steps of *P. anserina*’s life cycle as shown by a schematic representation (upper panel) and the corresponding light microphotographs (lower panel).** *P. anserina’s* life cycle begins with the germination of an ascospore (A) that gives rise to a haploid mycelium (B). After three days of growth, both male gametes (spermatia, B, top) and female gametes (ascogonia, B, bottom) are formed. Because most of the ascospores carry two different and sexually compatible nuclei (*mat*+ and *mat*-mating types) *P. anserina* strains are self-fertile (pseudo-homothallism). Before fertilization occurs, ascogonia can mature into protoperithecia by recruiting protective maternal hyphae to shelter the ascogonial cell. A pheromone/receptor signaling system allows the ascogonia to recognize and fuse with spermatia of compatible mating type (heterothallism). Fertilization initiates the development of the fruiting body (perithecia, C) in which the dikaryotic mat+/mat-fertilized ascogonium forms. Further development leads to a three-celled hook-shaped structure called the crozier (D). The two parental nuclei in the middle cell of the crozier fuse (karyogamy, schematic representation D) to form a diploid nucleus, which then immediately undergoes meiosis. The four resulting haploid nuclei undergo mitosis. In most cases, ascospores are formed around 2 non-sister nuclei within the developing ascus. On rare occasions, two ascospores are formed around only one haploid nucleus each, leading to a five-ascospore ascus (E, photograph). Scale bar: 10 µm in (A-D); 200 µm in (E).

**S4 Fig. Morphological phenotype of** Δ***PaRid* and complemented** Δ***PaRid*:AS4-PaRid-HA strains during vegetative growth.** Strains were grown on M2 minimal medium for 6 days at 27°C. S: Wild-type strain. See Material and Methods section for details on the complemented Δ*PaRid:AS4:PaRid:HA* strain.

**S5 Fig. Vegetative growth rates at various sub-optimal temperatures and longevity assays.** For each genotype, growth was assayed on 3 independent cultures after 4 days of growth at the indicated temperatures. Each experiment was performed three times. For each genotype, longevity was measured on three independent cultures, issued from three individual ascospores, as described in [41].

**S6 Fig. Perithecia versus micro-perithecia development with respect to genetic backgrounds.** (**A**) Morphological comparison of perithecia obtained from wild-type crosses (*PaRid*), Δ*PaRid* crosses (Δ*PaRid*) and Δ*Smr1* crosses (Δ*Smr1*). Size and morphology of the Δ*PaRid* and Δ*Smr1* perithecia are alike. Scale bar: 250 µm. (**B**) Perithecia obtained in the indicated trikaryons on M2 medium after 5 days at 27°C. The Δ*mat* ; *PaRid*^*+*^ *mat*-; *PaRid*^*+*^ *mat*+ trikaryons form typical fully developed perithecia (left panels). By contrast, only blocked micro-perithecia are formed by the Δ*mat* ; Δ*PaRid mat*-; Δ*PaRid mat*+ trikaryons (right panels). (**C**) Heterozygous orientated crosses *PaRid*^*+*^ *mat*-× Δ*PaRid mat*+ after 5 days at 27°C. When the wild-type *PaRid*^*+*^ allele is present in the female gametes genome and the mutant Δ*PaRid* allele is present in the male gamete genome fully developed perithecia are formed, conversely when the mutant Δ*PaRid* allele is present in the female gametes and the wild-type *PaRid*^*+*^ allele is present in the male gamete, only blocked micro-perithecia are formed. Left panel, scale bar: 1 mm, right panel, scale bar: 500 µm.

**S7 Fig. Western immunoblot analysis of the HA-tagged PaRid proteins expressed from the various ectopic *PaRid* alleles.** Western-blot was performed as described in Material and Methods section and probed with an anti-HA antibody that specifically detects the HA-tagged proteins. The PaRid protein (85 kDa) was barely detectable when the expression of the ectopic alleles was driven from its native promoter (Δ*PaRid*:*PaRid*, Δ*PaRid*:*PaRid*^*C403S*^). However, it was readily produced when the expression of the ectopic alleles was under the control of a strong and constitutive promoter (Δ*PaRid*:AS4-*PaRid*, Δ*PaRid*:AS4-*PaRid*^C403S^). Complementation of the fertility defect was obtained by insertion of the wild type *PaRid*-HA allele or by insertion of the AS4-*PaRid*-HA only. Δ*PaRid* mutant strain (negative control), Δ*PaRid*:*PaRid* = Δ*PaRid* mutant strain harboring an ectopic wild type *PaRid-HA* allele (complemented strain), Δ*PaRid*:AS4-*PaRid* = Δ*PaRid* mutant strain harboring an ectopic AS4-*PaRid*-HA allele (complemented strain), Δ*PaRid*:*PaRid*^C403S^ = Δ*PaRid* mutant strain harboring an ectopic catalytically dead *PaRid*^C403S^*-*HA allele (non-complemented strain), Δ*PaRid*:AS4-*PaRid*^C403S^ = Δ*PaRid* mutant strain harboring an ectopic AS4-*PaRid*^C403S^-HA allele (non-complemented strain). See the material and method section for details on the alleles construction and features.

**S8 Fig. Phylogenetic analysis of *P. anserina* down-regulated CDS encoding transcription factors and their orthologs in *N. crassa, A. immersus* and *A. nidulans* when present, using a maximum likelihood tree.** If four out of 17 have orthologs in *A. nidulans, A. immersus* and *N. crassa* (Pa_1_17860, Pa_1_22930, Pa_2_740 and Pa_2_5020), Pa_4_1960 is the only TF of the set showing no orthologs into the genomes of these three species. Although Pa_1_18880 and Pa_7510 display orthologs either in *N. crassa* or in *N. crassa* and *A. nidulans*, their phylogenetic positions were ambiguous, suggesting some species specialization. See Table S6 for protein names. Only bootstrap values > 0.5 are indicated on the corresponding branches.

**S1 Table.** Primers used in this study.

**S2 Table.** Vegetative phenotypic analyses.

**S3 Table.** Sexual reproduction phenotypic analyses.

**S4 Table.** Grafting experiments.

**S5 Table.** List of DE CDS. DE DOWN: down-regulated CDS. DE UP: up-regulated CDS. DOWN FC max: list of CDS with maximum fold change higher than −5. UP FC max: list of CDS with maximum fold change higher than +5. A: target of mating-type transcription factors, activated; R: target of mating-type transcription factors, repressed; 0: CDS not controlled by neither FPR1 nor FMR1. ^d^ = the corresponding gene was previously deleted [48].

**S6 Table.** List of CDS encoding transcription factors found down-regulated in the Δ*PaRid* micro-perithecia. A = ascospores, P = perithecia. ^d^ = the corresponding gene was previously deleted [48].

